# Functionally linked amygdala and prefrontal cortical regions are innervated by both single and double projecting cholinergic neurons

**DOI:** 10.1101/2023.12.29.573623

**Authors:** Bence Barabás, Zsófia Reéb, Orsolya I. Papp, Norbert Hájos

**Affiliations:** HUN-REN Institute of Experimental Medicine, 1083, Budapest, Hungary; János Szentágothai School of Neurosciences, Semmelweis University, 1085, Budapest, Hungary; Doctoral School of Biology, Institute of Biology, ELTE Eötvös Loránd University, 1117 Budapest, Hungary; The Linda and Jack Gill Center for Molecular Bioscience, Indiana University Bloomington, 47405, Indiana, USA; Program in Neuroscience, Department of Psychological and Brain Sciences, Indiana University Bloomington, 47405, Indiana, USA

**Keywords:** Basal forebrain, amygdala, prefrontal cortex, VGLUT3, ChAT, VGAT, cholinergic

## Abstract

Cholinergic cells have been proposed to innervate simultaneously those cortical areas that are mutually interconnected with each other. To test this hypothesis, we investigated the cholinergic innervation of functionally linked amygdala and prefrontal cortical regions. First, using tracing experiments, we determined that cholinergic cells located in distinct basal forebrain (BF) areas projected to the different nuclei of the basolateral amygdala (BLA). Specifically, cholinergic cells in the ventral pallidum/substantia innominata (VP/SI) innervated the basal nucleus (BA), while the horizontal limb of the diagonal band of Broca (HDB) projected to its basomedial nucleus (BMA). In addition, cholinergic neurons in these two BF areas gave rise to overlapping innervation in the medial prefrontal cortex (mPFC), yet their axons segregated in the dorsal and ventral regions of the PFC. Using retrograde-anterograde viral tracing, we demonstrated that a portion of mPFC-projecting cholinergic neurons also innervated the BLA, especially the BA. By injecting retrograde tracers into the mPFC and BA, we found that 28% of retrogradely labeled cholinergic cells were double labeled, which typically located in the VP/SI. In addition, we found that vesicular glutamate transporter type 3 (VGLUT3)-expressing neurons within the VP/SI were also cholinergic and projected to the mPFC and BA, implicating that a part of the cholinergic afferents may release glutamate. In contrast, we uncovered that GABA is unlikely to be a co-transmitter molecule in HDB and VP/SI cholinergic neurons in adult mice. The dual innervation strategy, i.e., the existence of cholinergic cell populations with single as well as simultaneous projections to the BLA and mPFC, provides the possibility for both synchronous and independent control of the operation in these cortical areas, a structural arrangement that may maximize computational support for functionally linked regions. The presence of VGLUT3 in a portion of cholinergic afferents suggests more complex functional effects of cholinergic system in cortical structures.

## Introduction

Recognizing threats is essential for animal survival. Therefore, it is not surprising that many brain circuits contribute to the detection of dangerous situations, leading to the elevation of attention and promoting affective brain state linked to the threat (Davis, 1997; Izquierdo et al., 2016; Ledoux, 2000; Tovote et al., 2015). This danger-triggered mental state helps generating the most appropriate behavioral responses, aiming to avoid, or at least, reduce any potential harm as well as forming memory that can guide future avoidance of similar threat. These complex brain processes are regulated at different levels in the nervous system. Top-down control is provided by cortical circuits located in the basolateral amygdala (BLA) and medial prefrontal cortex (mPFC)(Burgos-Robles et al., 2009; Herry et al., 2008; Little & Carter, 2013; Mcdonald et al., 1996; Sotres-Bayon et al., 2012; Weiskrantz, 1956). These two structures can be parceled based on their connectivity and role playing e.g., in the control of negative emotional states (Lacroix et al., 2000; Ledoux, 2000). Both the basal nucleus (BA) and the basomedial nucleus (BMA) of the BLA are reciprocally connected with the mPFC in a subregion-specific manner. BA forms connections with the prelimbic (PL) and anterior cingulate (ACC) cortices (Cassell et al., 1986; Gabbott et al., 2005; Vertes, 2004), whereas the BMA is interconnected with the infralimbic (IL) cortex (Hurley et al., 1991; Petrovich et al., 1996). The former functional unit has been shown to play a role in regulating high fear state and the related memory formation/storage (Kim et al., 2016a; Maren et al., 1996; Sierra-Mercado et al., 2011; Tye et al., 2011), while the latter one controls low fear state and is involved in extinction learning (Adhikari et al., 2015; Bloodgood et al., 2018; Lingawi et al., 2019).

As a general principle, cortical function is efficiently and rapidly affected by subcortical inputs in a brain state-dependent manner. One of the subcortical afferent systems contributing critically to cortical network operation originates from the basal forebrain (BF), a heterogeneous structure located in the medial-ventral part of the brain (Hasselmo & Sarter, 2011; Mesulam et al., 1983; Solari & Hangya, 2018; Zaborszky, 2002). In this brain region, there are three neuron types: cholinergic, GABAergic, and glutamatergic cells that are known to project to cortical areas (Farr et al., 1999; Goldbach et al., 1998; Pascual et al., 2004). Recent studies have established that GABA is often a co-transmitter molecule in cholinergic cells, implying complex effects on their postsynaptic neurons (Saunders et al., 2015; Takács et al., 2018). Interestingly, cholinergic cells in the nucleus basalis of Meynert of the BF have been proposed to project to cortical regions, e.g., to frontal and posterior cortical areas, that are functionally interconnected with each other (Gombkoto et al., 2021; Zaborszky et al., 1997; 2015). These results suggest that BF cholinergic inputs may orchestrate activity in functionally related cortical areas, promoting interaction between regions and ultimately, enhancing the neural computation. However, this attractive hypothesis has not been fully verified.

Previous studies have revealed that the BLA and mPFC receive cholinergic inputs primarily from the two main parts of the BF, the horizontal limb of the diagonal band (HDB) and ventral pallidum/ substantia innominata (VP/SI), respectively (Bloem et al., 2014; Carlsen et al., 1985; Mayo et al., 1984; Nagai et al., 1982). Whether the mPFC and BLA networks can be simultaneously or differentially regulated by cholinergic afferents conveying salient information has yet to be determined. It is a particularly important question as several studies have shown a role for the cholinergic system in fear related processes, including acquisition and memory recall (Crimmins et al., 2023; Jiang et al., 2016; Knox, 2016; Mineur et al., 2022; Power et al., 2003).

In this study we aimed to reveal the structural basis of BF cholinergic control over the interconnected BLA-mPFC fear circuits. Using retrograde and anterograde tracing techniques, together with immunocytochemistry, we mapped the projection areas of BF cholinergic neurons, first within the entire amygdala and prefrontal cortical regions (PFC), then specifically focusing on the innervation of the basolateral amygdala (BLA) and the medial prefrontal cortex (mPFC). We found that cholinergic neurons in the VP/SI preferentially target the BA, the dorsal part of the PFC (dPFC) and mPFC. In contrast, HDB cholinergic cells innervate predominantly amygdala regions surrounding the BA, such as BMA and the ventral part of the PFC (vPFC) together with the mPFC. Importantly, our findings demonstrate that a portion of cholinergic cells within the HDB and VP/SI simultaneously project to both the BLA and mPFC and the ratio of dual projecting cholinergic neurons can reach up to 30% within the VP/SI. Additionally, we found that vesicular glutamate transporter 3 (VGLUT3)-expressing neurons in the VP/SI form largely overlapping population with cholinergic cells, sharing similar projection patterns towards the BLA and mPFC. VGLUT3 was typically found in axon terminals located in the BA, but not in other BA-surrounding regions. On the other hand, vesicular GABA transporter (VGAT) might be expressed in cholinergic neurons of the VP/SI and HDB during development, but not in adult mice.

## Materials and methods

### Animals

In this study, C57Bl6 wild type mice, ChAT-IRES-Cre mice (www.jax.org, #006410), VGAT-IRES-Cre mice (www.jax.org, #016962), and the offspring of VGAT-Cre mice crossed with an Ai6 reporter mouse line (CAG-LSL-ZsGreen1, www.jax.org, #007906), as well as the offspring of BAC-VGLUT3-iCre mice (Slc17a8-iCre, www.jax.org, #018147) crossed with an Ai14 reporter mouse line (CAG-LSL-tdTomato, www.jax.or, #007914) were used. All experiments were performed in male and female mice 5-16 weeks of age. All experiments were approved by the Committee of the Scientific Ethics of Animal Research (22.1/360/3/2011) and all procedures involving animals were performed according to methods approved by Hungarian legislation (1998 XXVIII. section 243/1998, renewed in 40/2013.) and institutional guidelines of ethical code. All procedures complied with the European Convention for the Protection of Vertebrate Animals used for Experimental and Other Scientific Purposes (Directive 86/609/CEE and modified according to the Directives 2010/63/EU). All effort was taken to minimize animal suffering and the number of animals used.

### Retrograde tracing

Wild type mice were anaesthetized with 125 mg/kg ketamine and 5 mg/kg xylazine and mounted to a stereotaxic frame. Eyes were coated with corneal gel (Recugel Ophthalmic Gel), and body temperature was maintained with small animal heating pads (Supertech Instruments, Pecs, Hungary). The skin was removed from the skull, so the cranial sutures were clearly visible. The mouse head was set horizontally based on the level of the bregma and lambda. Anterior-posterior and medio-lateral coordinates were measured from the bregma. The skull was carefully drilled with a dental bur (Foredom), then dorso-ventral coordinates were measured from the level of the dura mater. To label BF neurons projecting to the amygdala region, we used Cholera Toxin B (CTB, 0.5 mg/99.5 µl distilled water, List Biological Laboratories) or Fast Blue (FB, 5% in in 0.9% saline, Polysciences), which were iontophoretically injected (2/2 s on/off duty cycle, 2 µA pulses, for 5 min for CTB, 2/2 s on/off duty cycle, 5 µA pulses, for 7-10 min for FB) into the lateral amygdala (LA) (AP/ML/DV coordinates, –1.7/3.4/3.6 mm), BA (AP/ML/DV coordinates, –1.5/3.2/4.2 mm) or BMA (AP/ML/DV coordinates, –1.7/2.8/4.6 mm) unilaterally using a Drummond Recording Nanoject II (Drummond Scientific). To label BF neurons projecting to the mPFC, FluoroGold (FG, 2% in 0.9% saline, Fluorochrome) was injected iontophoretically (2/2 s on/off duty cycle, 2 µA pulses, for 5 min; AP/ML/DV coordinates, 2.2, 1.9, 1.5/0.3/1 mm).

These retrograde tracers were applied to the brain via a glass pipette (ID = 0.530 mm ± 25 μm, OD 1.14 mm, World Precision Instruments) which was filled first with 50% glycerol (tap water: glycerol, 1:1) and then by the tracer. After 3-5 days following the injections, animals were anesthetized again with ketamine/xylazine mixture as above and perfused under deep anesthesia first with saline (0.9%) and then with 4% paraformaldehyde dissolved in 0.1 M phosphate buffer (PB, pH: 7.4, 100 mL/animal, Sigma-Aldrich). Brains were then removed and cut into 50-80 µm-thick coronal sections with a Vibratome (VT1000S, Leica).

### Anterograde tracing

To analyze the axonal arborization of BF cholinergic neurons projecting to the amygdala and the PFC, we injected 100-100 nl of AAV5.Ef1a.DIO.eYFP (gift from Karl Deisseroth; Addgene viral prep # 27056-AAV5, RRID: Addgene_27056) to the HDB (AP/ML/DV coordinates, 0.5/0.7/5.0 mm) and VP/SI (AP/ML/DV coordinates, 0.5/1.5/4.4 mm) within the same ChAT-Cre animals. To separately investigate the cholinergic projections of these BF nuclei, we injected 30 nl of AAV8.CAG.Flex.GFP virus (gift from Dr. Ed Boyden, UNC Vector Core) to the HDB (AP/ML/DV coordinates, 0.5/0.7/5.0 mm) or VP/SI (AP/ML/DV coordinates, 0.5/1.5/4.4 mm) into different ChAT-Cre mice. To investigate the glutamatergic projections of the VP/SI, we injected 30 nl of AAV8.CAG.Flex.GFP virus to the VP/SI of VGLUT3-Cre mice. Virus was injected with a Nanoject III Programmable Nanoliter Injector (Drummond Scientific). In this surgery, the glass pipettes were used as in retrograde tracing, but instead of glycerol, mineral oil (Sigma-Aldrich) was filled to the pipettes first, followed by introducing the virus. Injection speed was set to 3 nl/s until a clear change in the meniscus (mineral oil–virus border) was observed, then speed was set to 1 nl/s. After 4-5 weeks following the surgery, animals were anaesthetized and perfused the same way as described for retrograde tracing. Following tissue processing, we compared the normalized axonal fluorescent intensity (utilizing NIS elements AR 5.3) – obtained by anterograde virus tracing (HDB, VP/SI) – within the BLA and mPFC.

### Retrograde-anterograde virus tracing

To investigate the dual projection of BF cholinergic cells to the BLA and mPFC, we injected 3×100 nl AAV5.Ef1a.DIO.eYFP to the mPFC (AP/ML/DV coordinates 2.2, 1.9, 1.5/0.3/1 mm) unilaterally into ChAT-Cre mice. The injection method and parameters were consistent with those used for anterograde tracing. This approach allowed retrograde labeling of BF cholinergic neurons, expressing eYFP in their cell bodies as well as in their remote axons after 6-8 weeks.

### Immunostaining

Fixed brain sections were washed 3 times for 10 minutes in 0.1M PB, then a blocking solution (10% NDS, Normal Donkey Serum, 0.5% Triton-x 100 (10%), 0.1M PB) was applied for 30 minutes. The sections were incubated in a solution containing primary antibodies in addition to 2% NDS, 0.5% Triton-x 100 and 0.1M sodium azide dissolved in 0.1M PB for 1 day at room temperature, followed by 2 hours of incubation in the secondary antibody containing solution (1% NDS, 0.1M PB). Guinea pig anti-FG primary antibody (1:5000, Protos Biotech) and Alexa 488– or Cy3-conjugated donkey anti-guinea pig (1:500, Jackson ImmunoResearch) or goat anti-CTB primary antibody (1:20,000, List Biological Laboratory) and Cy3-conjugated donkey anti-goat (1:500, Jackson ImmunoResearch) were used to visualize FG– or CTB-containing BF neurons. Retrograde tracer Fast Blue, however, required no additional immunostaining. Rabbit anti-ChAT primary antibody (1:5000, SYSY) and Alexa 647– or Cy3-donkey anti-rabbit (1:500, Jackson ImmunoResearch) secondary antibody were used for visualizing ChAT+ neurons. Goat anti-GFP primary antibody (1:1000, Frontier Institute Co. Ltd) and Alexa 488-conjugated donkey anti-goat (1:500, Jackson ImmunoResearch) were used to enhance the visualization of virus-labelled axons.

To identify the neurochemical content of axon terminals in the amygdala, rabbit anti-ChAT (1:5000, SYSY) and rabbit anti-VAChT (vesicular acetylcholine transporter) primary antibody (1:1000, Frontier Institute Co. Ltd) and A488-conjugated donkey anti-rabbit secondary antibody (1:500, Jackson ImmunoResearch); goat anti-VGLUT3 primary antibody (1:1000, Frontier Institute Co. Ltd) and Cy3-conjugated donkey anti-goat secondary antibody (1:500, Jackson); guinea-pig anti-VGAT primary antibody (1:1000, Frontier Institute Co. Ltd) and Cy5-conjugated donkey anti-guinea pig secondary antibody (1:500, Jackson) were used. After the incubations, the sections were rinsed in 0.1M PB, and mounted on slides in Vectashield (Vector Laboratories).

### Confocal microscopy

Multicolor large images from different brain regions were taken with a C2 confocal laser scanning microscope (Nikon Europe, Amsterdam, The Netherlands) using 10x (Plan Fluor 10x, N.A. 0.3, xy: 1.25 μm/pixel) and 20x objectives (CFI Super Plan Fluor 20X, N.A. 0.45, xy: 0.58 μm/pixel). Multichannel images at a high resolution were acquired in channel series mode.

This way quantitative analysis and localization maps of retrogradely labeled BF cells could be performed on the same images. To quantify the neurochemical content of retrogradely labeled BF neurons, the NIS-Elements software was used. Then, large images were exported to Neurolucida software (10.53 software, MBF Bioscience), where the position of retrogradely labeled neurons as well as the brain structures and landmarks (ventral border of the slice, anterior commissure, midline) were indicated. This process was repeated at multiple coronal planes of the BF (0.62, 0.14, and –0.34 mm from bregma) in all injected animals. The drawings from the corresponding coronal planes were aligned and mapped onto the appropriate coronal section taken from the mouse brain atlas (Paxinos et al., 2004) using Adobe Photoshop (version 3.0, Adobe Inc.).

To reveal the neurochemical content of axon terminals in the amygdala region, confocal images were taken from each nuclei/subnuclei using a C2 microscope with a 60x objective (CFI Plan Apo VC60X Oil objective, N.A. 1.40; z (n=50) step size: 0.13 μm, xy: 0.08 μm/pixel). High resolution fluorescent images (2048×2048 pixel) from the injection sites in the HDB and VP/SI as well as amygdala region were taken using a 4x objective (C2 confocal laser scanning microscope, Plan Fluor 4x, N.A 0.13, xy: 1.54 μm/pixel). To quantify the VGAT content of BF cholinergic neurons, images were acquired in the HDB and VP/SI with a C2 confocal laser scanning microscope using a 10x objective (Plan Fluor 10x, N.A. 0.3, z (n=10) step size: 25.88 μm, xy: 1.25 μm/pixel).

### Axon arborization and injection parameter maps

To determine the proportion of the labeled axons in each amygdala nucleus and prefrontal cortical area, we imported the confocal images into the Adobe Photoshop, quantified the pixels representing axons in the region of interest and compared it to the total number of pixels from the axonal arborization within the amygdala or prefrontal cortical regions. To determine the borders of the amygdala nuclei based on soma distributions, we mapped the amygdala region at the coronal plains using NeuN staining (1:1000, Millipore) or we used reconstructed maps from the mouse brain atlas. In the case of the PFC, we only used edited pictures from the mouse brain atlas. All maps containing injection location and spread were made based on the mouse brain atlas (Paxinos et al., 2004). Data are presented as mean ± SD.

## Results

### Cholinergic innervation of the amygdala region by two basal forebrain areas, the HDB and VP/SI

First, we aimed to investigate the overall cholinergic innervation of the amygdala region given rise by the HDB and VP/SI. To this end, we injected a high volume of AAV5.Ef1a.DIO.eYFP, a Cre-dependent adeno-associated virus vector (AAV) into the HDB and VP/SI of ChAT-Cre mice (100-100 nl to each area, Fig. 1A, n=3, Suppl. Fig. 1A). After analyzing the cholinergic projection patterns in multiple coronal planes within the amygdala region, we found that the BA received the strongest cholinergic innervation – based on normalized fluorescence intensity – from these two BF areas (Fig. 1B-C). Cholinergic innervation extended also to the BMA, the medial division of the central amygdala (CeM), the anterior– and posteromedial cortical amygdaloid nucleus (ACo, PMCo), the anterolateral portion of the amygdalohippocampal area (AHiAL), and the piriform cortex (Pir) with comparable innervation levels among them (Fig. 1B-C). Conversely, the anterior and posterior sections of the medial amygdala (MeA, MeP), the lateral nucleus of the amygdala (LA), the lateral division of the central amygdala (CeL), and the posteromedial part of the amygdalohippocampal area (AHiPM) received only sparse cholinergic innervation (Fig. 1B-C). Taken together, these findings demonstrate that the amygdala region, particularly the BA and BMA, receives robust cholinergic innervation from the HDB and VP/SI.

**Fig. 1.**
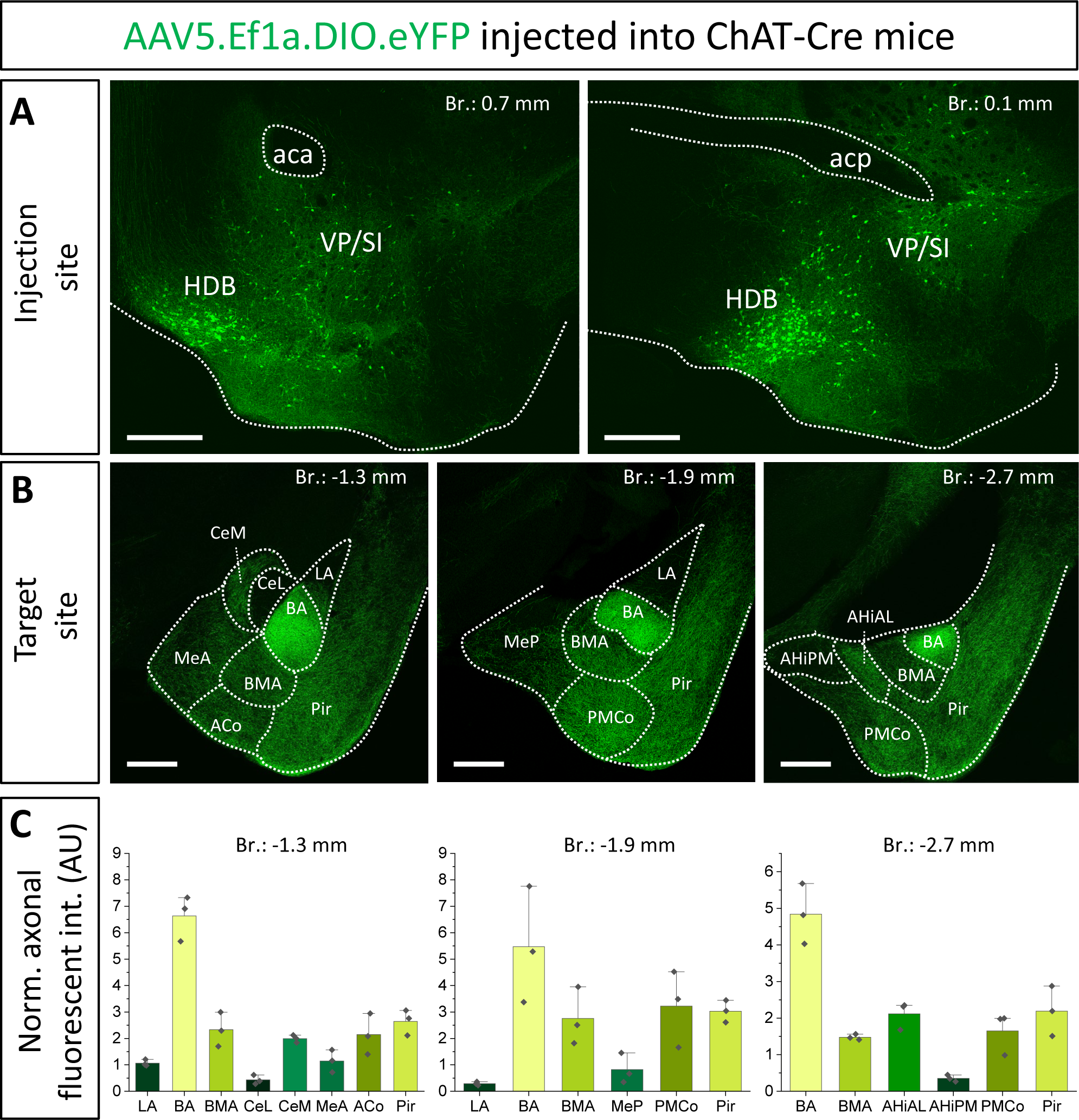
Cholinergic innervation of the amygdala region by two basal forebrain nuclei, the HDB and VP/SI. (A_1-2_) Example injection sites in the HDB and VP/SI in ChAT-Cre mice, 100 nl of AAV5.Ef1a.DIO.eYFP was injected to each area. Scale bar: 500 µm. (B_1-3_) Coronal images taken at different amygdala planes showing distribution of cholinergic projections from the HDB and VP/SI. Scale bar: 500 µm. (C) Normalized axonal fluorescent intensity (AU) graph comparing the abundance of cholinergic innervation within the amygdala region originating from the HDB and VP/SI. Aca, anterior commissure, anterior part; ACo, anterior cortical amygdaloid nucleus; acp, anterior commissure, posterior part; AHiAL, amygdalohippocampal area, anterolateral part; AHiPM, amygdalohippocampal area, posteromedial part; BA, basal amygdala; BMA, basomedial amygdala; CeL, central amygdala, lateral division; CeM, central amygdala, medial division; HDB, nucleus of the horizontal limb of the diagonal band; LA, lateral amygdala; MeA, medial amygdala, anterior part; MeP, medial amygdala, posterior part; Pir, piriform cortex; PMCo, posteromedial cortical amygdaloid nucleus; SI, substantia innominata; VP, ventral pallidum.

### Localization of BLA-projecting cholinergic neurons shows separation within the BF

To distinguish the sources of cholinergic innervation invading the BLA between the VP/SI and HDB, retrograde tracer Fast Blue (FB) or Cholera Toxin B subunit (CTB) was injected into the three distinct nuclei of the BLA: LA (Fig. 1A_1_, n=2), BA (Fig. 2A_2_, n=4) and BMA (Fig. 2A_3_, n=2). In coronal sections obtained from these injected mice, we determined the location of the somata of BLA-projecting neurons in the BF (Fig. 2B_1_). We found that BA-projecting BF neurons were located primarily in the VP/SI region (based on the mouse brain atlas (Paxinos et al., 2004)). In contrast, BMA-projecting BF neurons were found dominantly in the HDB (but also partially in the lateral preoptic area (LPO), median preoptic nucleus (MnPO), medial preoptic area (MPA)), forming a separate neuronal population, the location of which overlapped minimally with BA-projecting BF neurons (Fig. 2C_1_-C_3_). Injecting a retrograde tracer into the LA resulted in only a small number of labeled neurons located in the HDB (Fig. 2C_1_), a finding consistent with the low number cholinergic fibers observed in anterograde tracing (Fig. 1B-C). After performing immunostaining against choline acetyltransferase (ChAT), the enzyme, responsible for the acetylcholine synthesis (Jope, 1979), we evaluated the cholinergic content of the amygdala-projecting BF neurons. We found that more than 60% of BA-projecting BF neurons were cholinergic (Fig. 2B_1-3_ 130/212; n= 4 mice), while this ratio was around 25% (24/91; n= 2 mice) among BMA-projecting BF neurons. Altogether, these results show that neurons in the VP/SI preferentially innervate the BA, while those neurons located in the HDB project predominantly to the BMA and to a lesser extent to the LA. The difference in the ratio of BF cholinergic neurons innervating the distinct amygdala nuclei is in accord with our anterograde labeling, showing more profound presence of cholinergic fibers in the BA in comparison with the surrounding areas (Fig. 1B-C).

**Fig. 2.**
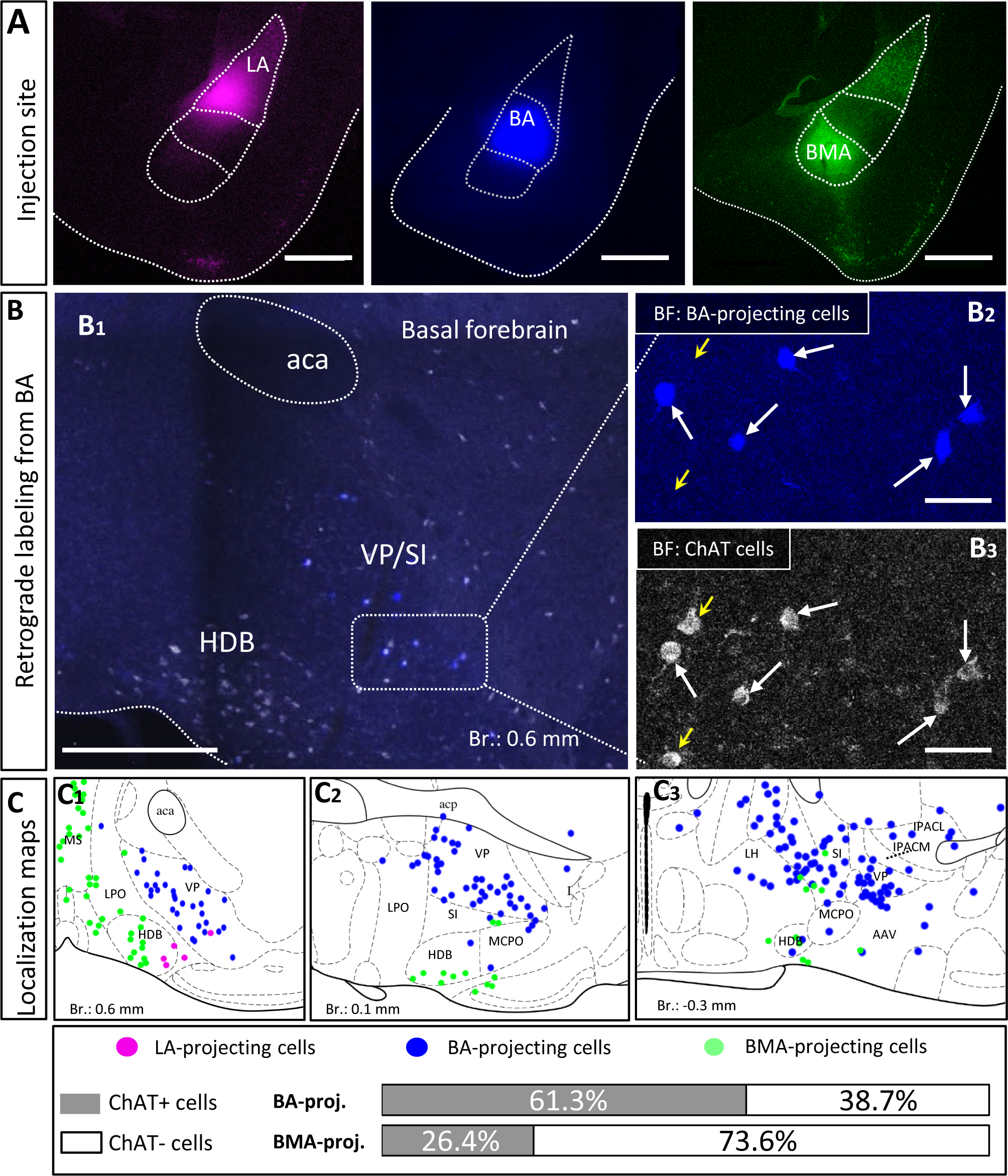
Distribution and cholinergic content of retrogradely labeled BLA-projecting BF neurons. (A) Pseudo-colored images showing the injection sites in different BLA regions. (A_1_) lateral amygdala, (LA); (A_2_) basal amygdala, (BA); (A_3_) basomedial amygdala, (BMA). Scale bar: 500 µm. (B_1_) An example large image taken from the basal forebrain (Br.: 0.62 mm, BF) showing BA-projecting BF cells in blue, and ChAT-immunoreactive cells in white. Scale bar: 500 µm. (B_2-3_) 20x magnification images taken from the BF showing expression of ChAT (choline acetyltransferase, B_3_) in BA-projecting BF cells (B_2_). Small yellow arrows indicate non-retrogradely labeled ChAT-expressing cells. Big white arrows indicate ChAT-positive BA-projecting cells. Scale bar: 50 µm. (C_1-3_) Positions of retrogradely labeled BLA-projecting BF neurons in different coronal planes of the BF. 61.3±7.08% of BA-projecting BF cells expressed ChAT (130/212; n= 4 mice), while the presence of ChAT was observed in 26.37 ±7.71% of BMA-projecting BF cells (24/91; n= 2 mice). Purple: LA-projecting cells, Blue: BA-projecting cells, Green: BMA-projecting cells. Aca, anterior commissure, anterior part; HDB, nucleus of the horizontal limb of the diagonal band; IPACL, interstitial nucleus of the posterior limb of the anterior commissure, lateral part; IPACM, interstitial nucleus of the posterior limb of the anterior commissure, medial part; LH, lateral hypothalamic area; LPO, lateral preoptic area; MCPO, magnocellular preoptic nucleus; MS, medial septal nucleus; SI, substantia innominata; VP, ventral pallidum.

### Cholinergic cells from the HDB and VP/SI project to the amygdala region in a mutually exclusive manner

To confirm that cholinergic neurons in separate BF regions innervate different amygdala nuclei, we injected a small amount (30 nl) of AAV8.CAG.Flex.GFP either into the HDB (Fig. 3A, n=3, Suppl. Fig. 1B) or into the VP/SI (Fig. 3D, n=4, Suppl. Fig. 1C) of ChAT-Cre mice. This approach enabled us to specifically investigate the cholinergic axonal projections originated from the HDB and VP/SI within the amygdala region. In line with our retrograde tracing data, we observed that ChAT+ neurons in the HDB projected mostly to the BMA and other surrounding areas, such as the MeA, ACo, dorsal endopiriform nucleus (DEn), Pir, but largely avoiding the BA (Fig. 3B, C, G). Although, there were some labeled axons in the LA, this connection appeared to be weak compared to other neighboring fields, an observation, which is in accord with the results shown in Fig. 1B-C, and Fig. 2C_1_. On the other hand, ChAT+ neurons in the VP/SI almost exclusively innervated the BA, with marginal projections into the CeM, BMA and Pir (Fig. 3E-G). These results show that cholinergic fibers from the HDB and VP/SI parcel the amygdala region in a mutually exclusive manner.

**Fig. 3.**
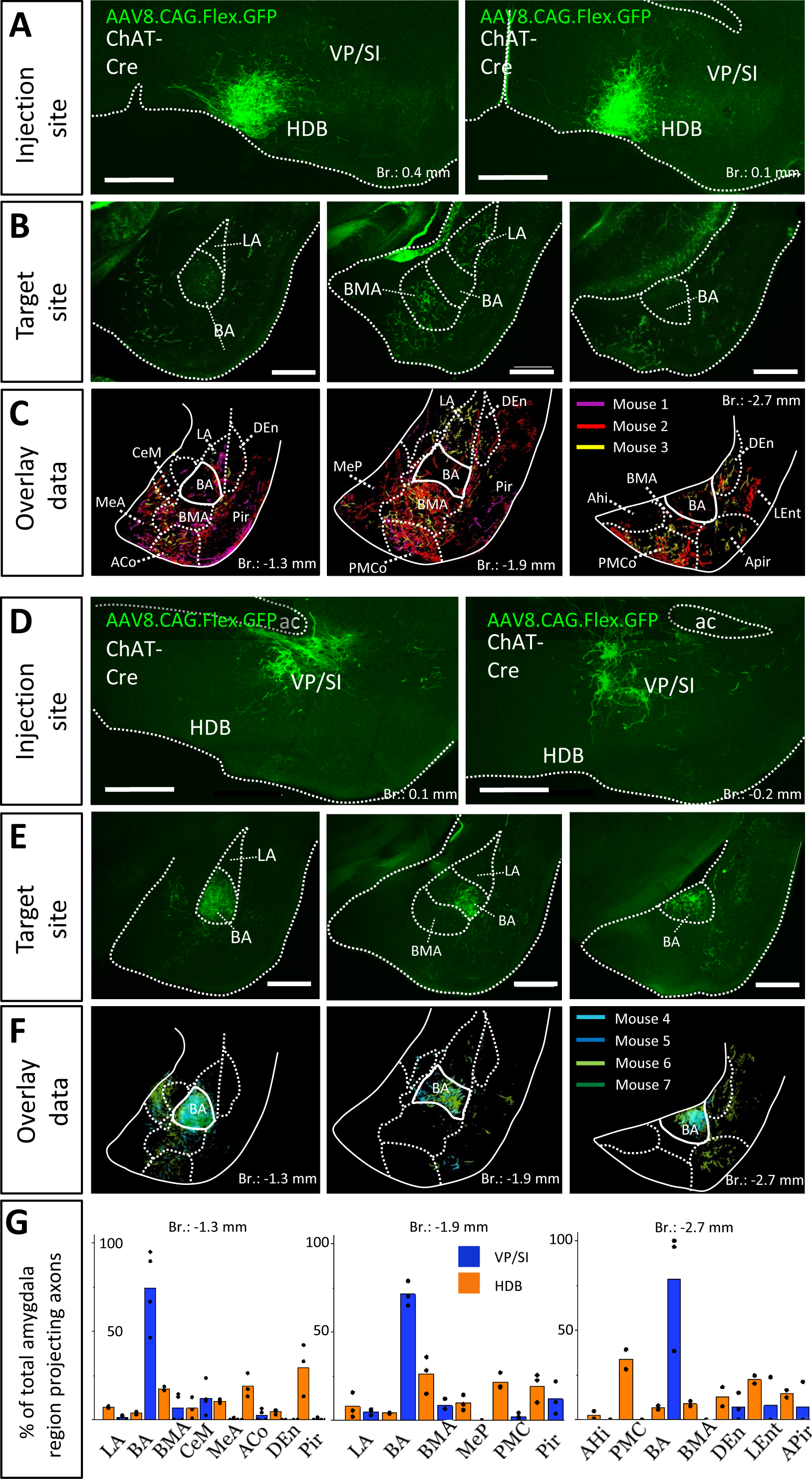
Distribution of anterogradely labeled cholinergic projections in the amygdala region from the HDB or VP/SI. (A) Example injection sites in the HDB in ChAT-Cre mice. Small volume (30nl) of AAV8.CAG.flex.GFP was injected to the target area. Scale bar: 500 µm. (B) Coronal images taken at different amygdala planes showing cholinergic projections from the HDB. Scale bar: 500 µm. (C) Overlaid representative cholinergic projections from the HDB. Axons visualized in different animals (n=3) are shown in different colors. Borders of distinct amygdala regions were drawn based on NeuN staining. (D) Example injection sites in the VP/SI in ChAT-Cre mice. 30nl AAV8.CAG.flex.GFP was injected to the target area. Scale bar: 500 µm. (E) Coronal images taken at different amygdala planes showing cholinergic projections from the VP/SI. Scale bar: 500 µm. (F) Overlaid representative cholinergic projections from the VP/SI. Axons revealed in different animals (n=4) are shown in distinct colors. The borders of distinct amygdala regions are drawn based on NeuN staining. (G) Percentage of amygdala region-projecting HDB or VP/SI cholinergic axons in each nucleus/region. ACo, anterior cortical amygdaloid nucleus; AHi, amygdalohippocampal area; APir, amygdalopiriform transition area; BA, basal amygdala; BMA, basomedial amygdala; CeM, central amygdala, medial division; DEn, dorsal endopiriform nucleus; LA, lateral amygdala; LEnt, lateral entorhinal cortex; MeA, medial amygdala, anterior part; Mep, medial amygdala, posterior part; Pir, piriform cortex; PMCo, posteromedial cortical amygdaloid nucleus.

### Cholinergic inputs from the HDB and VP/SI overlap in the mPFC, with VP/SI showing a preference for dorsal and HDB for ventral innervation of the PFC

Previous studies uncovered that the mPFC and BLA are reciprocally interconnected cortical regions playing a role in similar cognitive processes (Arruda-Carvalho & Clem, 2015; Burgos-Robles et al., 2009; Herry et al., 2008; Little & Carter, 2013; Sotres-Bayon et al., 2012; Weiskrantz, 1956). Therefore, we analyzed the cholinergic projections from the HDB and VP/SI towards the PFC in the same mice as we did for the innervation of the amygdala region (Figs 1 and 3). Upon analyzing the cholinergic projection patterns towards the PFC (Fig. 4A, same injections as shown in Fig. 1), we observed that cholinergic fibers covered the cingulate cortex area 1 and 2 (Cg1, Cg2), as well as the PL and IL cortices and the medial orbital cortex (MO) in a similar manner (Fig. 4B-C). Additionally, these fibers exhibited a slightly weaker innervation to the secondary motor cortex (M2) (Fig. 4B-C). Based on small injections targeted separately into the HDB or VP/SI (injections shown in Fig. 3), we found that both BF areas projected to the mPFC, namely the ACC/Cg1, PL and IL cortices. However, the HDB tended to project to the vPFC, including the MO, lateral orbital cortex (LO), ventral orbital cortex (VO), dorsal peduncular cortex (DP), Cg2 (Fig. 5A, B, E), while the VP/SI innervated the dPFC (including the M2, Cg1 and PL; Fig. 5C-E). The most conspicuous differences were found at the level of 2.4 mm from the bregma, where the HDB predominantly innervated the MO, while the VP/SI had prominent projections to the PL, Cg1 and M2 (Fig. 5B, D). At the bregma level of 1.1 mm, there was a similar exclusive projection pattern: cholinergic cells in the HDB innervated the Cg2, but not the M2, while cholinergic cells in the VP/SI gave rise to projections to the M2, but not to the Cg2 (Fig. 5B, D). Altogether, these results show that cholinergic innervation originated from the HDB and VP/SI terminates in the differential parts of the PFC and amygdala regions. Namely, ChAT+ neurons in the HDB send axons to the vPFC and mPFC as well as to the most amygdala areas, apart from the BA, while ChAT+ neurons in the VP/SI prefer to terminate in the dPFC, mPFC and BA.

**Fig. 4.**
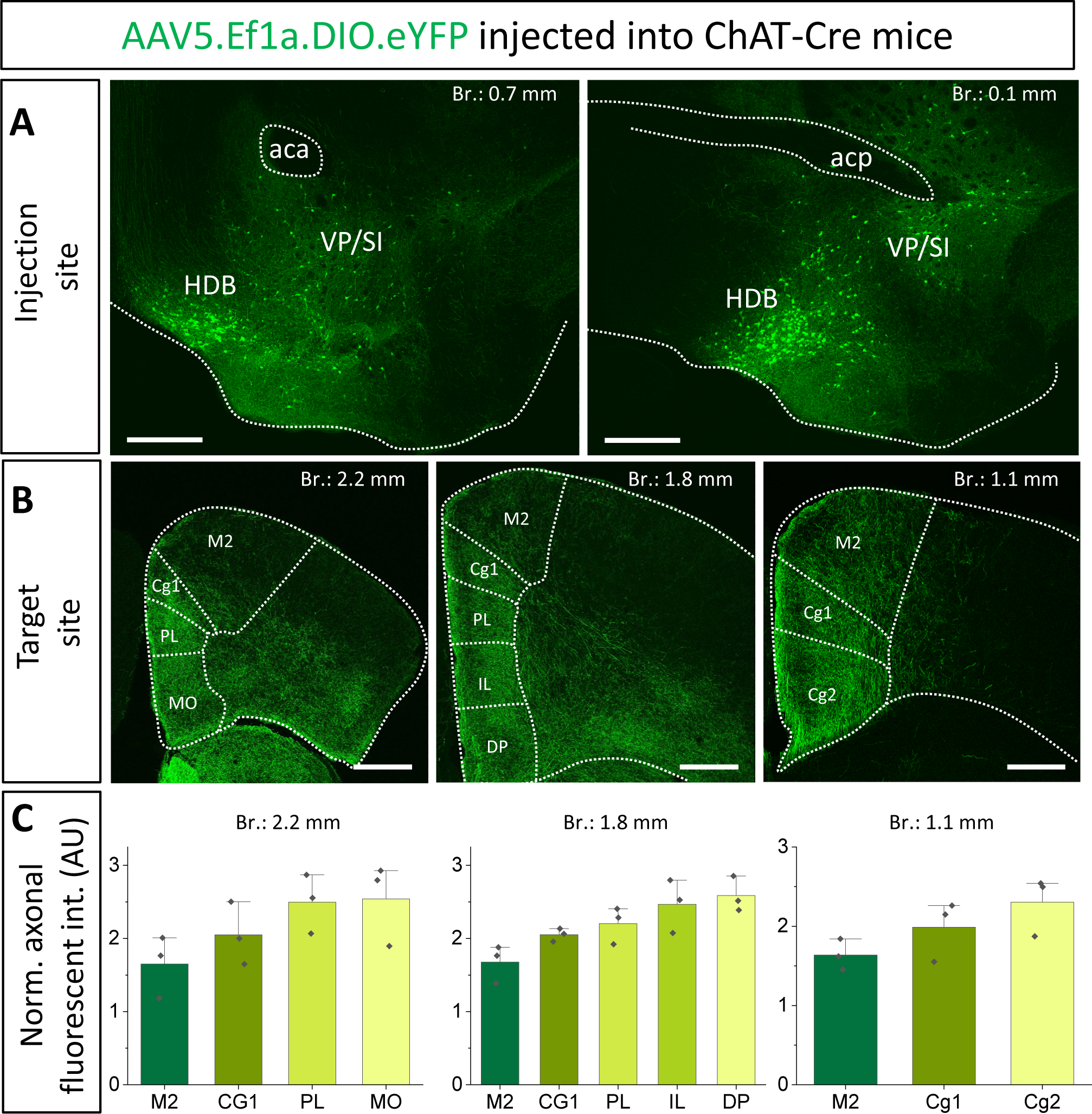
Cholinergic innervation of the prefrontal cortex by the HDB and VP/SI. (A_1-2_) Example injection sites in the HDB and VP/SI in ChAT-Cre mice. Large volume (100nl) of AAV5.Ef1a.DIO.eYFP was injected to each area. Scale bar: 500 µm. Same images as shown in Fig. 1A. (B_1-3_) Coronal images taken at different prefrontal cortical planes showing cholinergic projections from the HDB and VP/SI. Scale bar: 500 µm. (C) Normalized axonal fluorescent intensity (AU) graph comparing the abundance of cholinergic innervation within the PFC region originating from the HDB an VP/SI. Aca, anterior commissure, anterior part; acp, anterior commissure, posterior part; Cg1, cingulate cortex, area 1; Cg2, cingulate cortex, area 2; DP, dorsal peduncular cortex; IL, infralimbic cortex; LO, lateral orbital cortex; M1, primary motor cortex; M2, secondary motor cortex; MO, medial orbital cortex; PL, prelimbic cortex; VO, ventral orbital cortex.

**Fig. 5.**
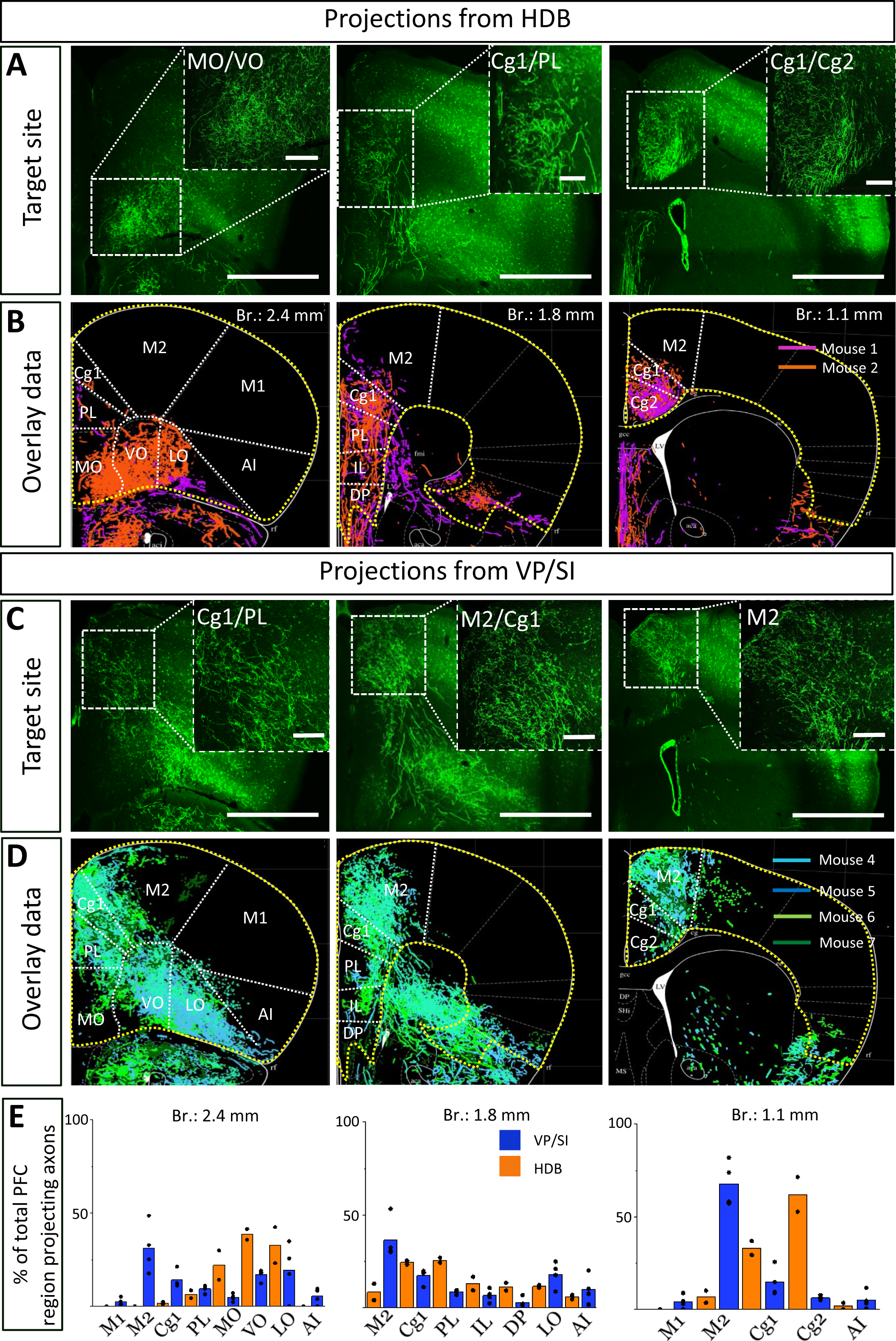
Cholinergic afferents from the HDB and VP/SI highly overlap in the mPFC but show separation in the dorsal and ventral parts of the PFC. (A, C) Images taken at different prefrontal cortical planes showing cholinergic projections from the HDB (A) and VP/SI (C) (injection sites presented in Fig. 3). Scale bar: 1000 µm. (B, D) Overlaid representative cholinergic projections from the HDB (B) or VP/SI (D). Axons represented in different colors were obtained from different animals (n=2-4). Maps of the PFC were modified from the mouse brain atlas (Paxinos et. al., 2004). (E) Percentage of PFC-projecting cholinergic axons in each cortical region from the HDB or VP/SI. AI, agranular insular cortex; Cg1, cingulate cortex, area 1; Cg2, cingulate cortex, area 2; DP, dorsal peduncular cortex; IL, infralimbic cortex; LO, lateral orbital cortex; M1, primary motor cortex; M2, secondary motor cortex; MO, medial orbital cortex; PFC, prefrontal cortex; PL, prelimbic cortex; VO, ventral orbital cortex.

### Significant portion of BF cholinergic neurons exhibit dual projections to both the mPFC and BLA

The next question we asked was whether the same BF neurons send axonal collaterals into the mPFC and BLA, or separate populations of neurons innervate these two brain structures. To reveal the logic underlying the cholinergic control of these regions, first we performed retrograde-anterograde virus tracing in ChAT-Cre mice by injecting AAV5.Ef1a.DIO.eYFP into the mPFC (Fig. 6B, n=5). This approach reveals whether neurons projecting to a given area have axonal collaterals in other brain regions, too. Thus, AAV injection into the mPFC may visualize axons in the BLA if a portion of BF cholinergic neurons simultaneously project to these regions. Our retrograde-anterograde virus tracing demonstrated the presence of ChAT+ neurons in the BF (Fig. 6C, Suppl. Fig. 2) that innervate both the mPFC (Fig. 6B) and BLA (Fig. 6D-E). The somata of retrogradely labeled ChAT+ cells were found in both the HDB and VP/SI along the BF (Suppl. Fig. 2), revealing the area where the dual projecting cholinergic neurons were located. Based on overlaid images (Fig 6E), extracted from 5 animals, we found that BA received a substantial cholinergic innervation from dual projecting neurons, with the strongest innervation found at the level of –1.5 mm from the bregma (Fig. 6E-F). BMA also received axon collaterals from the dual projection, while there were barely any fibers in the LA (Fig. 6E-F). Our results provide evidence that the mPFC-BLA circuit receives dual cholinergic innervation from both the HDB and VP/SI.

**Fig. 6.**
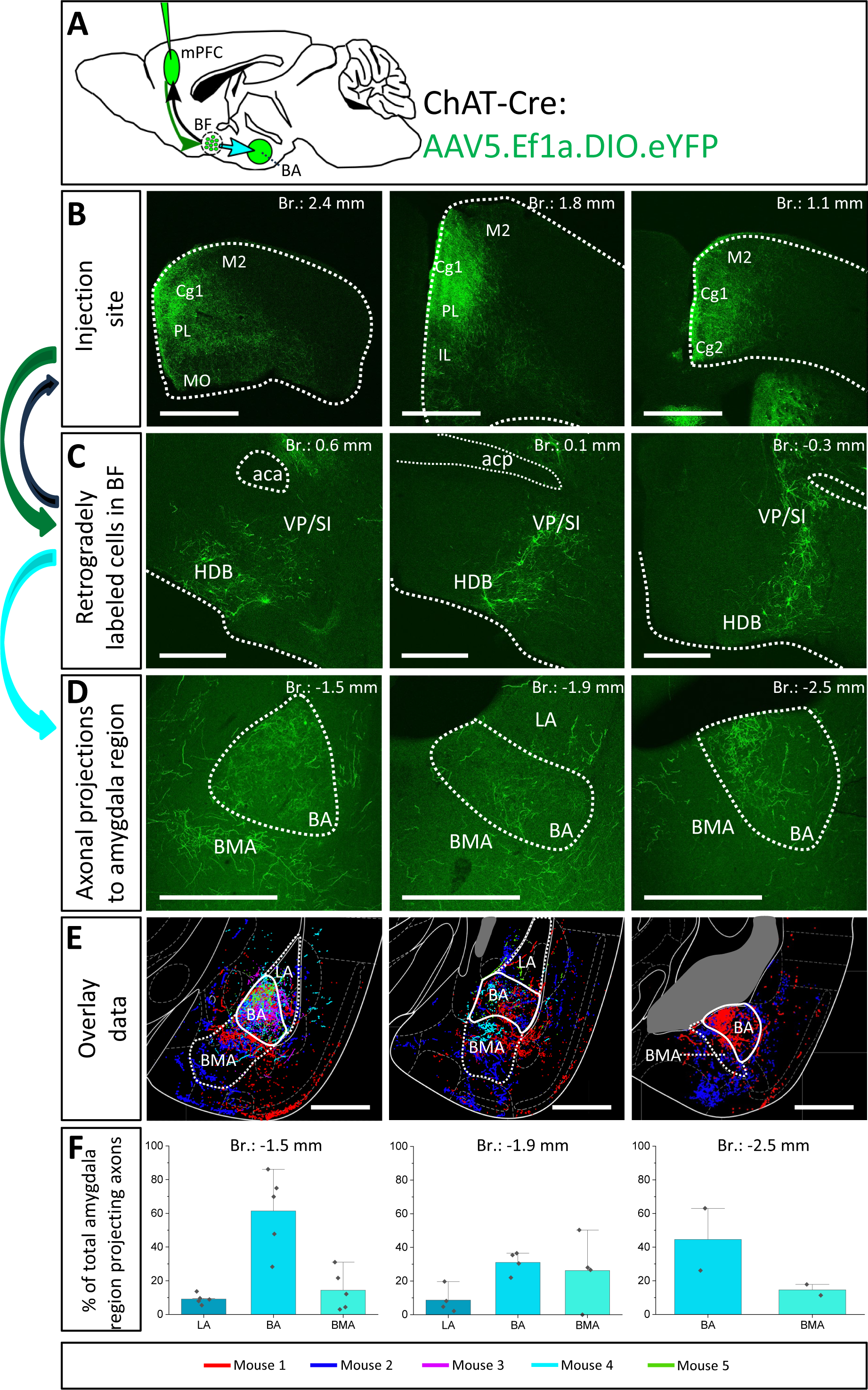
Retrograde-anterograde virus tracing demonstrates the presence of a cholinergic neuronal population with dual projection towards the mPFC and BLA. (A) Schematic representation of retrograde-anterograde Cre-dependent axonal labeling. AAV5.Ef1a.DIO.eYFP was injected into the mPFC (3 x 100 nl) in ChAT-Cre mice (n = 5). Green arrow shows retrograde spread of the Cre-dependent virus retrogradely to BF cholinergic cells. Black arrow indicates anterograde labeling of axons towards the injection site. Cyan arrow shows anterograde labeling of axons towards the amygdala. (B) Coronal images taken from the injection sites in the mPFC showing cholinergic fibers from retrogradely labeled cholinergic BF cells. Scale bar: 1000 µm. (C) Coronal images of retrogradely labeled cell bodies of processes of cholinergic neurons in the BF. Scale bar: 500 µm. (D) Fluorescent images of BLA-projecting axon collaterals in the of those BF cholinergic neurons, which also innervated the mPFC. Scale bar: 500 µm. (E) Overlaid representative axonal projections in the BLA originating from dual projecting cholinergic BF cells. Axons revealed in different animals (n=5) are shown in distinct colors. Scale bar: 500 µm. (F) Percentage of BLA-projecting cholinergic axons in the BA, BMA and LA. BA, basal amygdala; BMA, basomedial amygdala; Cg1, cingulate cortex, area 1; Cg2, cingulate cortex, area 2; IL, infralimbic cortex; LA, lateral amygdala, M1, primary motor cortex; M2, secondary motor cortex; MO, medial orbital cortex; PL, prelimbic cortex; HDB, nucleus of the horizontal limb of the diagonal band; SI, substantia innominata; VP, ventral pallidum.

To further validate the observation that cholinergic neurons simultaneously innervate the mPFC and BLA, we injected different retrograde tracers into the mPFC and BA (Fig. 7A-C, Suppl. Fig. 3, n=3), areas receiving the strongest innervation from cholinergic cells with dual projections (Fig. 6). In line with our previous tracing results, retrogradely labeled neurons from the mPFC were located both in HDB and VP/SI (Fig. 7D-E_1_, E_3_), while BA-projecting cholinergic cells were mostly found in the VP/SI (Fig. 7D-E_1_, E_3_). Importantly, we identified a population of dual-projecting cells in the VP/SI (Fig. 7F), accounting for 30.07±12.5% (Fig. 7G, 109/355, n=3) of the retrogradely labeled cells, the vast majority of which (92.7%) was immunopositive for ChAT (Fig. 7E_4_, G, 101/355, n=3), leaving a small portion (2.2±1.2%, Fig. 7G, 8/355, n=3) of dual-projecting cells immunonegative for ChAT. These results confirm that BF cholinergic cells can regulate the interaction between the mPFC and BLA simultaneously and/or independently.

**Fig. 7.**
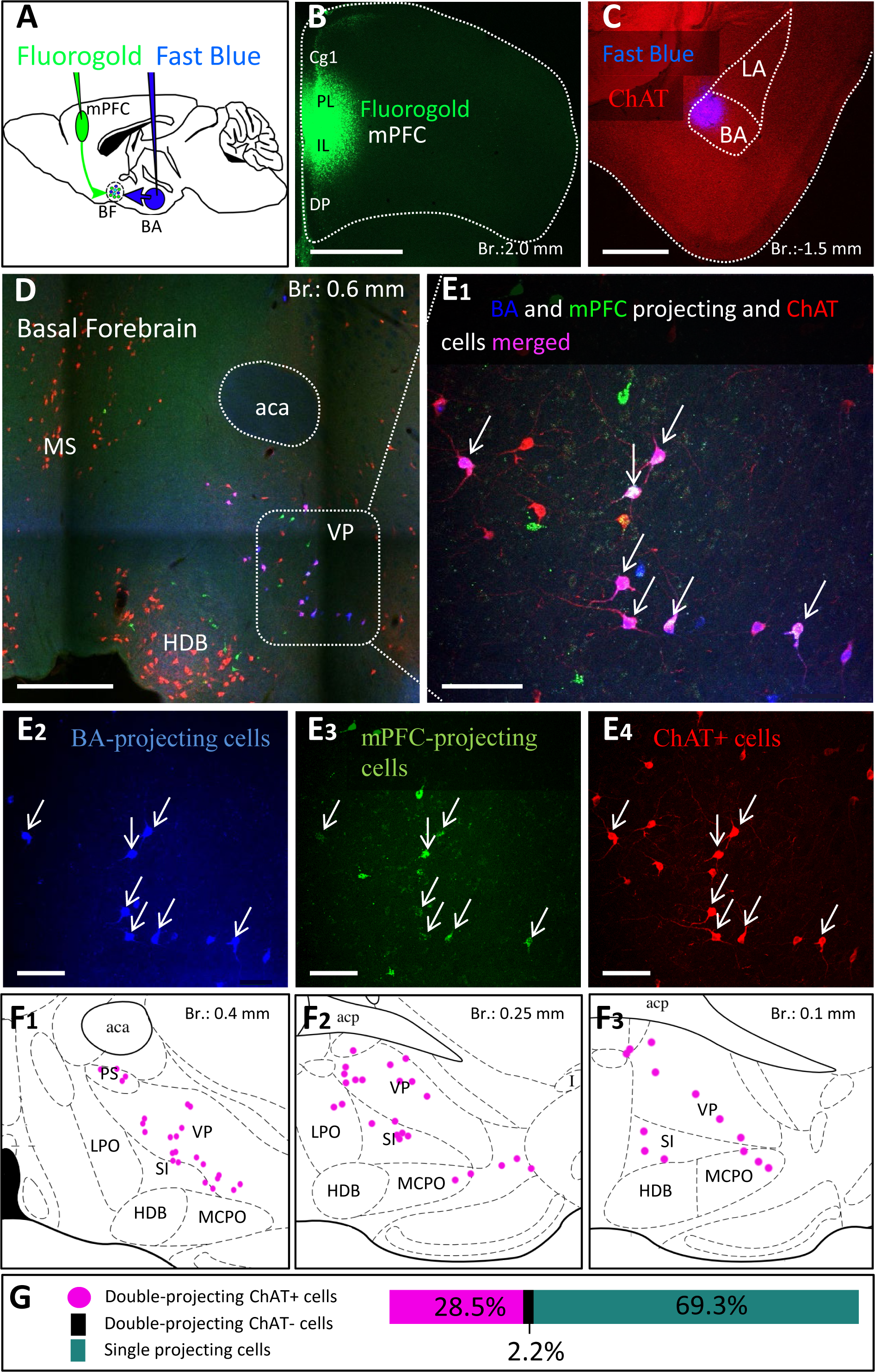
Double-retrograde labeling of BF neurons that project to the mPFC and BA. (A) Experimental design: iontophoretic injection of Fluorogold into the mPFC and Fast Blue to the BA. (B) Example coronal image taken at the injection site in the mPFC showing Fluorogold labeling in the PL and IL regions. Scale bar: 1000 µm. (C) Example coronal image taken from the same animal showing the injection site of Fast Blue labeling in the BA. Scale bar: 500 µm. (D-E_1-4_) 10x and 20x magnification coronal images at a BF plane showing BA-projecting (E_2_), mPFC-projecting (E_3_) and ChAT-immunopositive (E_4_) cells in the VP/SI. White arrows indicate double-labeled cholinergic neurons. Scale bar: (D) 500 µm, (E1-4) 100 µm. (F_1-3_) Localization of double-projecting cholinergic neurons shown in purple in different planes of the BF. (G) Bar graph showing that 28.5±12.48% (n =3, 101/355) of retrogradely labeled neuros were double-projecting and ChAT-positive, while only 2.2±1.22% (n=3, 8/355) were double-projecting and ChAT-negative. 69.3±11.5% of retrogradely labeled neurons were single-projecting (n=3, 246/355). Purple: double-projecting ChAT-positive neurons; Black: double-projecting ChAT-negative neurons; Green: single-projecting neurons. Aca, anterior commissure, anterior part; BA, basal amygdala; Cg1, cingulate cortex, area 1; DP, dorsal peduncular cortex; HDB, nucleus of the horizontal limb of the diagonal band; IL, infralimbic cortex; LA, lateral amygdala; LPO, lateral preoptic area; MCPO, magnocellular preoptic nucleus; MS, medial septal nucleus; PL, prelimbic cortex; PS, parastrial nucleus; SI, substantia innominata; VP, ventral pallidum.

### ChAT and VGLUT3 cells form highly overlapping populations within the VP/SI

A previous study obtained in rats has uncovered (Poulin et al., 2006) that BF cholinergic neurons projecting to the BLA express VGLUT3. To clarify whether amygdala-projecting cholinergic neurons also contain VGLUT3 in mice, we revealed the neurons expressing VGLUT3 by crossing VGLUT3-Cre and Ai14 reporter mice. In offspring, we performed immunostaining against ChAT that allowed us to quantify the number of cholinergic neurons in the HDB and VP/SI that expressed VGLUT3, too. We observed that 71.19% (168/236, n=2) of cholinergic neurons in the VP/SI expressed VLGUT3, whereas this ratio was 33.69% (94/279, n=2) in the HDB (Suppl. Fig. 4).

Knowing that most cholinergic cells also contain VGLUT3 in the VP/SI, we administered 30 nl AAV8.CAG.Flex.GFP into the VP/SI of VGLUT3-Cre mice to explore the projection patterns and cholinergic phenotype of VGLUT3+ neurons and their axons in the PFC and amygdala region (Fig. 8, Suppl. Fig. 1D).We found that 64% (Fig. 8A, 89/139, n=2) of VGLUT3+ neurons were also positive for ChAT at the injection site, and these cells exhibited similar projection patterns towards the amygdala (Fig. 8B) and PFC (Fig. 8D), mirroring our observations obtained in ChAT-Cre mice. The most prominent difference was that VGLUT3+ axons in the BA could not be observe in posterior amygdala planes (from Br.: –1.8mm to –2.9 mm), while fewer axons were found in the M2 cortex (Suppl. Fig. 5). However, more prominent projections were seen in the anterior insular cortex (AI, Fig. 8D, Suppl. Fig. 5). Additionally, ChAT immunostaining revealed that VGLUT3+ axons contained ChAT enzyme in both the BA (Fig. 8C) and mPFC (mainly Cg1, PL) (Fig. 8E), further supporting the observation that VGLUT3+ and ChAT+ neurons form overlapping populations in the VP/SI.

**Fig. 8.**
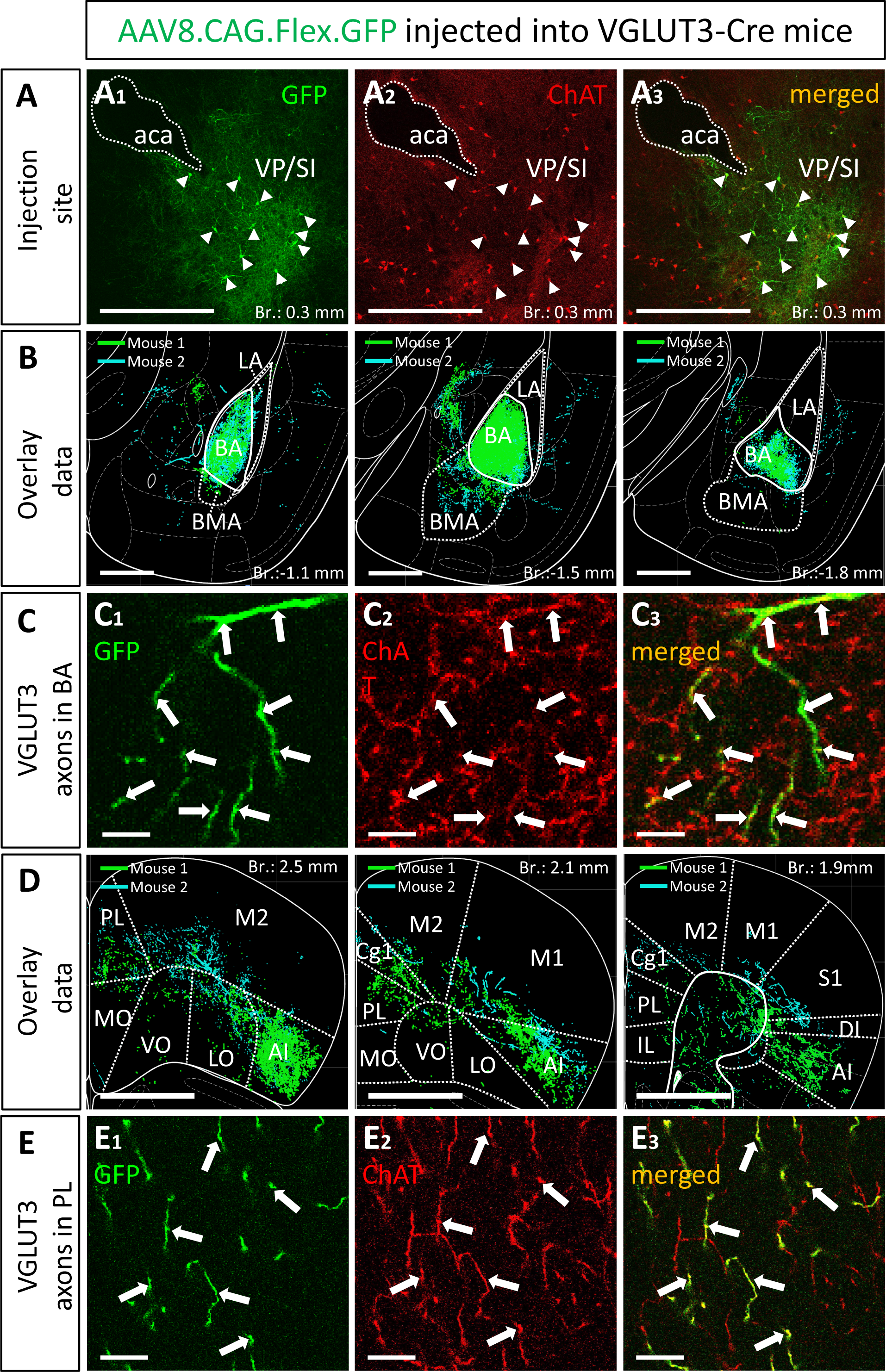
VGLUT3-expressing neurons within the VP/SI show ChAT immunoreactivity and project to the mPFC and BA. (A) Coronal image taken from the BF showing the injection site (A_1_) in the VP/SI in VGLUT3-Cre mice. Small amount (30nl) of AAV8.CAG.flex.GFP was injected to the target area. Cholinergic cells were visualized with ChAT immunostaining (A_2_) to reveal the cholinergic content of VGLUT3-expressing BF cells (A_3_) in the VP/SI. Scale bar: 500 µm. White arrowheads: double labeled neurons. (B) Overlaid representative glutamatergic projections in the BLA originating from VP/SI cells. Axons revealed in different animals (n=2) are shown in distinct colors. Scale bar: 500 µm. (C) 60x magnification fluorescent images showing VGLUT3-GFP (C_1_) and ChAT (C_2_) co-labeling (C_3_) within the BA. Scale bar: 5 µm. White arrows: double labeled axons. (D) Overlaid representative glutamatergic projections in the mPFC originating from VP/SI cells. Axons revealed in different animals (n=2) are shown in distinct colors. Scale bar: 1000 µm. (E) 60x magnification fluorescent images showing VGLUT3-GFP (E_1_) and ChAT (E_2_) co-labeling (E_3_) within the PL. Scale bar: 10 µm. White arrows: double labeled axons. Aca, anterior commissure, anterior part; AI, agranular insular cortex; BA, basal amygdala; BMA, basomedial amygdala; Cg1, cingulate cortex, area 1; IL, infralimbic cortex; LA, lateral amygdala, LO, lateral orbital cortex; M1, primary motor cortex; M2, secondary motor cortex; MO, medial orbital cortex; PL, prelimbic cortex; VO, ventral orbital cortex; S1, primary somatosensory cortex; DI, dysgranular insular cortex; SI, substantia innominata; VP, ventral pallidum.

Collectively, these data predict that most cholinergic axon terminals within the BA should also contain VGLUT3 immunolabeling, whereas the ratio of the double labeled axonal boutons for ChAT and VGLUT3 should be substantially less in other amygdala areas. In agreement with this hypothesis, we found that, among all amygdala areas, the BA had the highest ratio of axon terminals immunoreactive for both ChAT and VGLUT3 (Fig. 9A). Interestingly, this ratio was not homogeneous within this nucleus: its anterior part (Fig. 9A, C, 63.28%± 8.58, 324/512, n=3) contained twice as many double immunopositive axonal boutons than its posterior part (Fig. 9C, 29.67%± 3.51, 89/300, n=3). This observation might explain why fewer VGLUT3+ fibers were observed in caudal amygdala planes (Fig. 8B). As expected, in neighboring regions, the co-expression of ChAT and VGLUT3 was minimal (Fig. 9B, C). These results together suggest that cholinergic pathways from the VP/SI typically contain VGLUT3, while ChAT+ fibers located in other amygdala regions – originating primarily from the HDB – express this glutamate transporter at a much lower level. In addition, these observations further strengthen the conclusion that the BA and its surrounding regions receive cholinergic control from two distinct BF sources that differ in their VGLUT3 content.

**Fig. 9.**
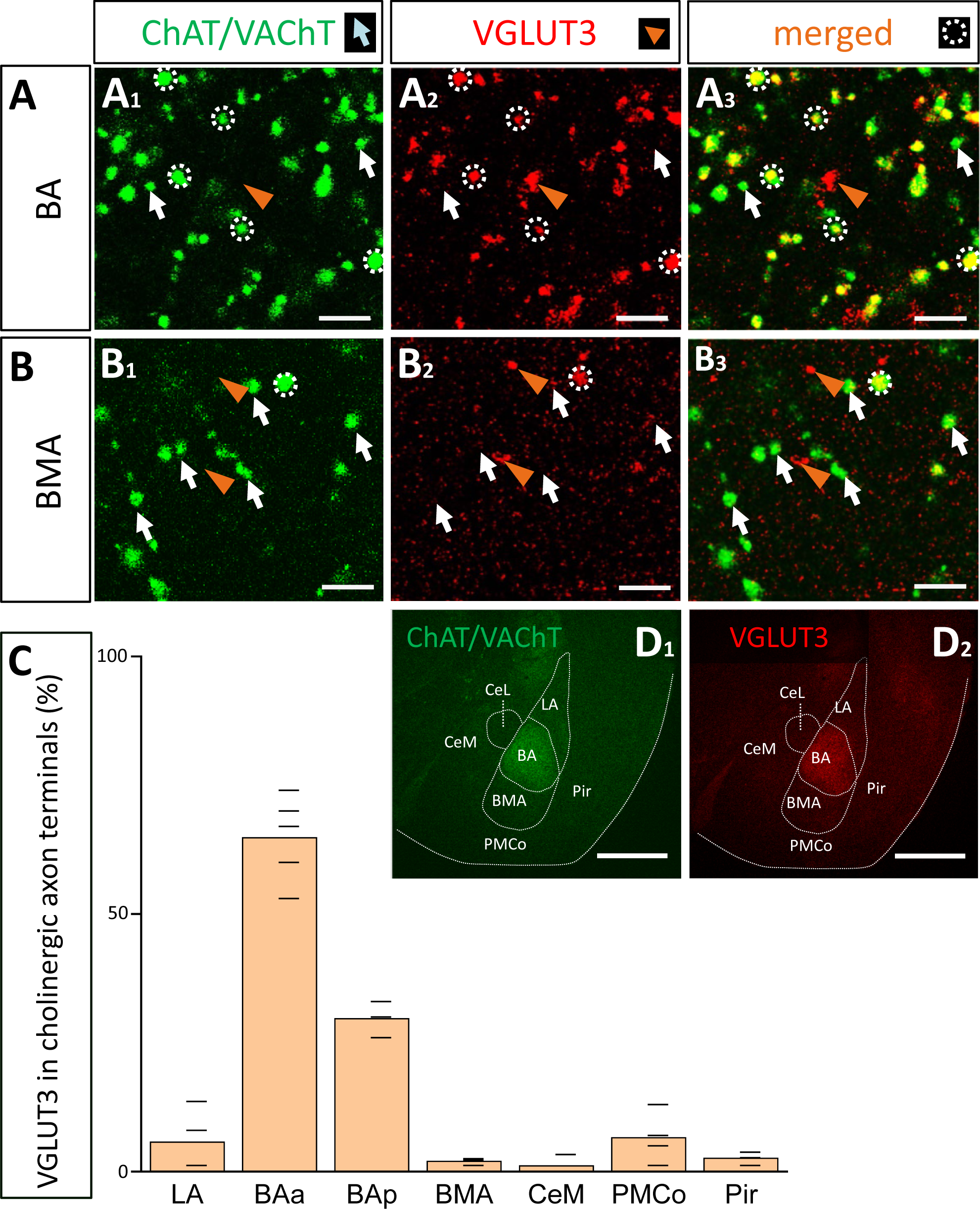
Cholinergic axon terminals in the BA co-express VGLUT3. (A) Cholinergic (A_1_; green. ChAT/VAChT) and glutamatergic axon terminals (A_2_; red. VGLUT3) revealed by immunostaining showing high co-occurrence in cholinergic boutons in the BA (A_3_). Scale bar: 3 µm. White arrows point to ChAT/VAChT-containing boutons. Orange arrowheads indicate VGLUT3-containing boutons. White dotted circles indicate double labeled i.e., cholinergic and VGLUT3-immunostained boutons. Scale bar: 3 µm. (B) Cholinergic (B_1_; green) and VGLUT3-containing axonal boutons (B_2_; red) form largely non-overlapping populations in the BMA. Scale bar: 3 µm. (C) Graph showing that the VGLUT3 content in cholinergic axon terminals is the highest in the anterior BA (BAa; 64.8 ± 8.35%; n=3, 332/512) and posterior BA (BAp; 29.67 ± 3.51%, n=3, 89/300), while the co-expression is lower in surrounding amygdala nuclei: LA (5.82 ± 6.33%, n=2, 17/292), BMA (2.38 ± 0.68%, n= 2, 6/252), PMCo (6.85 ± 4.93%, n=2, 24/350), Pir (2.57 ± 1.05%, n=2, 8/311), CeM (1.69 ± 1.92%, n=2, 3/177). (D) Images taken at low magnification from ChAT/VAChT (D_1_; green) and VGLUT3 (D_1_; red) immunostained sections showing high expression levels for these proteins in the BA, but not in surrounding amygdala regions. Scale bar: 500 µm. BAa, basal amygdala, anterior part; BAp, basal amygdala, posterior part; BMA, basomedial amygdala; CeM, central amygdala, medial division; LA, lateral amygdala; Pir, piriform cortex; PMCo, posteromedial cortical amygdaloid nucleus.

### GABAergic phenotype is not characteristic for cholinergic neurons in the HDB and VP/SI in adult mice

Previous studies convincingly demonstrated that GABA can be released from cholinergic axon terminals in the neocortex (Saunders et al., 2015), hippocampus (Takács et al., 2018) and striatum (Lozovaya et al., 2018). To assess whether cholinergic neurons in the HDB and VP/SI are also GABAergic, we visualized inhibitory cells in the BF by crossing VGAT-Cre with Ai9 reporter mice (CAG-LSL-ZsGreen1, Fig. 10A_1_, B_1_), whereas ChAT content of neurons was revealed by immunostaining (Fig. 10A_2_, B_2_). We found that 96.98% (546/563, n=3) of cholinergic neurons expressed the reporter protein under the control of VGAT promoter in the HDB, whereas this ratio was 58.8% (137/233, n=3) in the VP/SI (Fig. 10E, dark green). To confirm the GABAergic nature of cholinergic afferents in the BLA, we examined the VGAT immunoreactivity in cholinergic axon terminals. To our surprise, we observed no co-expression of VGAT and cholinergic markers (ChAT; VAChT) (Fig. 10F, 0.75%, 3/400, n=2) in axonal boutons at the amygdala level. To resolve this contradiction, we labeled BF GABAergic cells in adult VGAT-Cre mice by injecting AAV8.CAG.Flex.GFP vector simultaneously into the HDB and VP/SI (Fig. 10C_1_, D_1_). After 4 weeks following the injections, we performed immunostaining against ChAT (Fig. 10C_2_, D_2_) and counted the GFP content of cholinergic cells in the BF (Fig. 10C_3_, D_3_). Under these circumstances, we found that there were only a few neurons co-expressing ChAT and GFP both in the HDB (3.06%, 19/620, n=3) and VP/SI (1.19%, 3/253, n=3) (Fig. 10E, light green). These data are in line with the findings that cholinergic axon terminals do not contain VGAT immunolabeling in the amygdala (Fig. 10F). Thus, our results collectively show that in adult mice, GABA is not typically present in cholinergic neurons in the HDB and VP/SI.

**Fig. 10.**
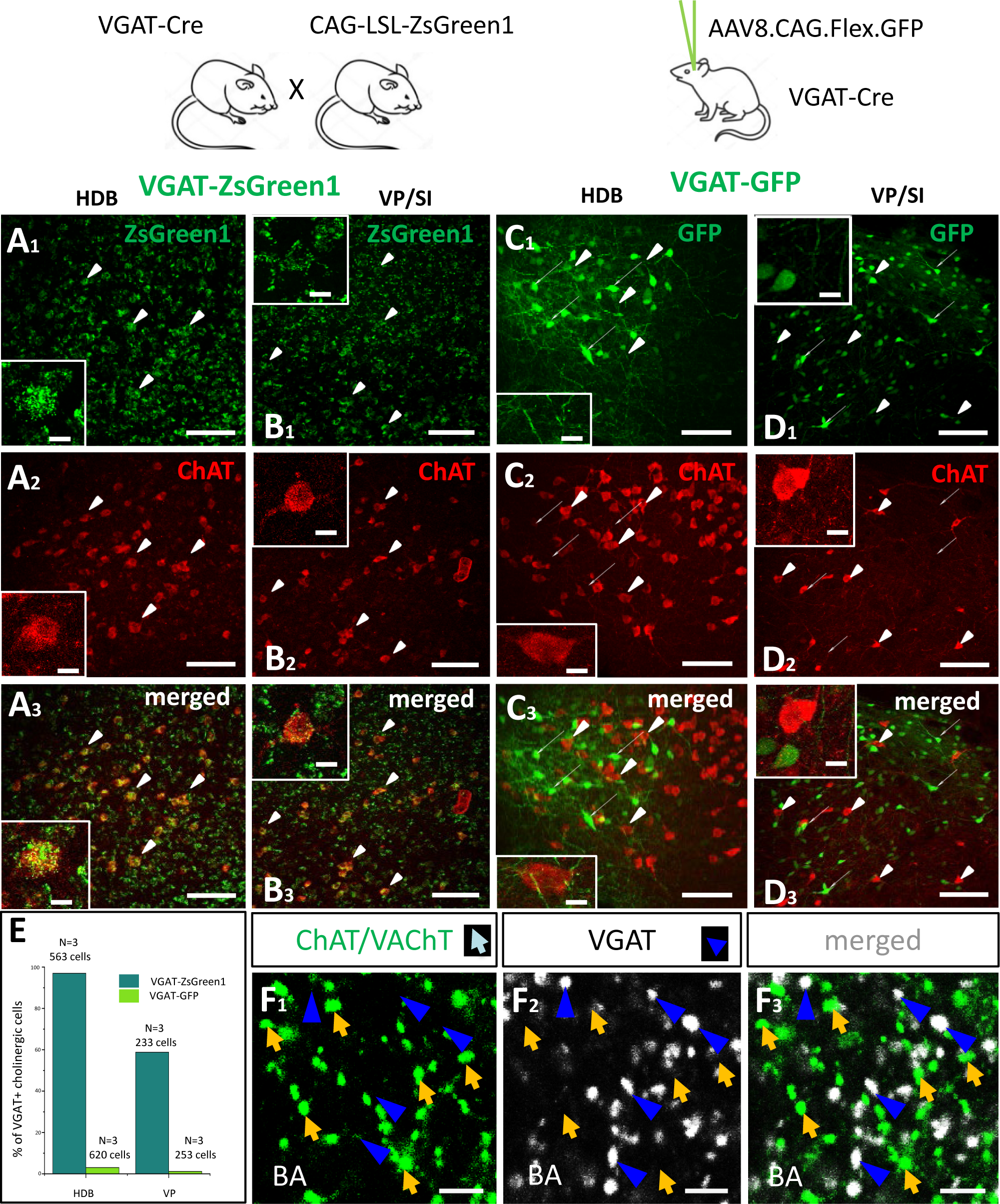
Cholinergic cells in the HDB and VP/SI do not express VGAT in adult mice. (A, B) Visualizing GABAergic cells in the HDB (A_1_) and VP/SI (B_1_) within the BF, in VGAT-ZsGreen1 offspring generated by breeding VGAT-IRES-Cre and CAG-LSL-ZsGreen1 mice. Scale bar: 50 µm. Insets: images taken at higher magnification of a GABAergic BF neuron containing ChAT immunolabeling. Scale bar: 5 µm. (C, D) Virus labeling of GABAergic cells in the HDB (C_1_) and VP/SI (D_1_) by injecting AAV8.CAG.Flex.GFP into VGAT-Cre mice. Scale bar: 50 µm. Insets: images taken at higher magnification showing GABAergic and cholinergic cells intermingled in the BF. Scale bar: 5 µm. (A_2_, B_2_, C_2_, D_2_) ChAT immunostaining revealing cholinergic cells in the BF. Scale bar: 50 µm. (A_3_, B_3_) Merged images showing co-expression of VGAT and ChAT both in the HDB and VP/SI in VGAT-ZsGreen1 mice. Scale bar: 50 µm. (C_3_, D_3_) Merged images showing that cholinergic cells in the HDB and VP/SI do not contain VGAT in AAV-injected VGAT-Cre mice. Scale bar: 50 µm. (E) Percentage of VGAT-expressing cholinergic cells in the HDB (Dark green, 96.7%, 546/563, n=3) and VP/SI (Dark green, 58.8%, 137/233, n=3) in VGAT-ZsGreen1 and AAV-injected VGAT-Cre (Light green, HDB: 3.06%, 19/620; VP/SI: 1.19%, 3/253, n=3) mice. (F_1-3_) Cholinergic (F_1_; green. ChAT/VAChT) and GABAergic (F_2_; white. VGAT) axon terminals showing non-overlapping populations in the BA (F_3_). Scale bar: 3 µm. Orange arrows show ChAT/VAChT-containing boutons. Dark blue arrowheads point to VGAT-immunoreactive boutons.

## Discussion

In this study we have investigated the cholinergic innervation of the reciprocally interconnected mPFC-BLA regions. Our main findings are the followings (Fig11). 1) Cholinergic cells located in two separate BF areas, the HDB and VP/SI innervate different parts of the BLA in a mutually exclusive manner. Although, they exhibit overlapping projections in the mPFC, the HDB tends to project more to the vPFC, whereas the VP/SI prefers to innervate the dPFC. 2) Importantly, using retrograde-anterograde virus tracing and double-retrograde tracing, we provide evidence that a significant portion of cholinergic cells can simultaneously innervate the BLA and mPFC. 3) Dual-projecting cells are intermingled with single-projecting neurons within the HDB and VP/SI. 4) Cholinergic and VGLUT3-containing BF cells form largely overlapping populations in the VP/SI and exhibit similar projection patterns toward the amygdala region and PFC. Moreover, VGLUT3+ axons originate primarily from the VP/SI and express ChAT in both the mPFC and BA. 5) VGLUT3 is much less typical for cholinergic cells and their axon terminals in the HDB and other amygdala areas. 6) Cholinergic cells in the HDB and VP/SI do not express VGAT in adult mice, thus, they unlikely have GABAergic phenotype (Fig. 11).

**Fig. 11.**
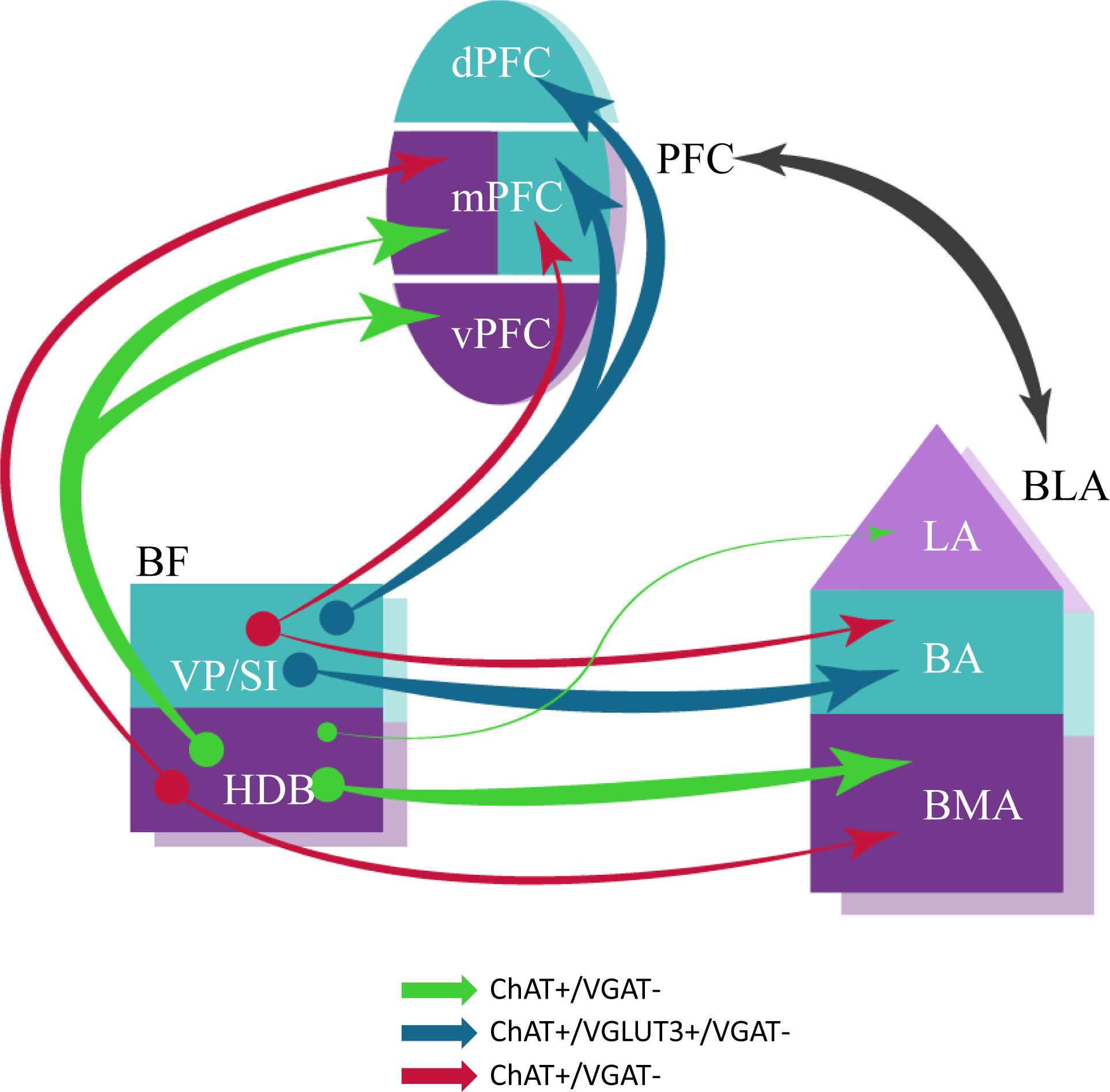
Schematic illustration of BF cholinergic inputs to the interconnected PFC-BLA networks. Both single– and dual-projecting BF cholinergic cells give rise to the innervation of the functionally linked PFC and BLA. This structural arrangement of cholinergic inputs can allow the maximalization of computational support for coordinated functioning of interconnected cortical regions. Anatomical and functional related brain regions are indicated in the same colors. The thickness of the arrows corresponds to the abundance of cholinergic innervation. The color of the arrows represents the neurochemical content of the projecting neurons. BA, basal amygdala; BLA, basolateral amygdala; BMA, basomedial amygdala; dPFC, dorsal part of the PFC; HDB, nucleus of the horizontal limb of the diagonal band; LA, lateral amygdala; mPFC, medial PFC; PFC, prefrontal cortex; VP/SI, ventral pallidum/substantia innominata; vPFC, ventral part of the PFC.

Our results are in a good agreement with previous observations showing that the majority of BF neurons (61%) located in the VP/SI innervating the BA express ChAT (Carlsen et al., 1985; Poulin et al., 2006). In addition, we provide evidence that the HDB innervates the BMA, yet the ratio of cholinergic neurons in this pathway is considerably lower (25%). The difference in the proportion of cholinergic neurons within the BF afferents terminating in the BA and BMA is also reflected in the abundance of cholinergic fibers in these amygdala nuclei (Fig. 1B-C).

In our study we found striking similarities in the distribution of BF afferents innervating the PFC to that observed earlier (Bloem et al., 2014). However, we substantially expanded these pervious findings by showing that the same BF regions distinctly projected to the amygdala region. Thus, beside the extensively studied nucleus basalis of Meynert (Crouse et al., 2020; Jiang et al., 2016; Kitt et al., 1987; Mallet et al., 1995), the VP/SI gives rise to a significant innervation to the BA, while the HDB provides inputs to additional amygdala areas albeit less intensively. We also observed cholinergic projection from the VP/SI to the CeM, in agreements with a previous study (Aitta-aho et al., 2018). Based on our tracing experiments, two parallel cholinergic innervation patterns can be outlined: 1) the VP/SI to the dPFC – mPFC and BA and 2) the HDB to the vPFC – mPFC and BMA/LA projections. While these parallel pathways overlap in the mPFC, they may have differential effects on circuit operation in subregions like PL and IL, potentially regulating opposing fear states (Burgos-Robles et al., 2009; Santini et al., 2008; Senn et al., 2014). Collectively, these neuronal circuits may play a crucial role in conveying salient information to the BLA and mPFC, thereby regulating functions associated with fear and reward processing, as well as social memory and interaction (Jiang et al., 2016; Kim et al., 2016; Lee & Kim, 2019; Okada et al., 2021; Robert et al., 2021). A recent study reported that the HDB is important in 1) predicting whether a perceptual outcome in a behavioral task was hit or miss and 2) encoding the omission of anticipated rewards, while the VP/SI can encode 1) nociceptive inputs and auditory stimuli and 2) learning related plasticity of auditory cues associated with punishment (Robert et al., 2021). These observations highlight the distinct functions for the two parallel cholinergic pathways.

Our retrograde-anterograde tracings not only confirmed the presence of dual-projecting cholinergic cell axons in the mPFC and BLA, but also defined the area in the BF where these neurons were located. To reveal the ratio of double-projecting cells, we used double retrograde tracing between the mPFC and BA. Based on these experiments, we found that ∼ 30% of VP/SI cholinergic cells project to both cortical structures. This observation is somewhat contradictory to earlier observations obtained in rats (Carlsen et al., 1985), where the authors barely found any double-labeled neurons between the BA and the neo– or allocortex. In our study, we noticed that different retrograde tracers or viruses had different success in labeling BF cells. For example, when we injected AAVretro.CAG.eGFP or AAVretro.CAG.tdTomato into the mPFC and/or BLA, we found no retrogradely labeled cells in the BF, but there were many retrogradely labeled cells in the other regions, including the thalamus, PFC and BLA. Similarly, using Alexa-conjugated CTB, we only found 1-3 retrogradely labeled BF cells per section, whereas using FG and Fast Blue with the same injection parameters, we observed 10-20 retrogradely labeled BF cells in a section. Carlsen et al. used Fast Blue combined with Nuclear Yellow, an approach that might have different success compared to our method, while they also injected into mostly different cortical regions as we did. Furthermore, injecting a small amount of tracers into brain areas, which have extensive rostro-caudal spread – such as the mPFC and amygdala – might not cover the axonal fields of BF cells in both areas, therefore, the double-projecting cholinergic cells may be missed. Meanwhile, larger injections have the risk to spread into neighboring regions, resulting in unwanted labeling. To overcome this issue, we applied multiple small injections within the mPFC and BA along the anterior-posterior axis. In line with our findings, Li et al., 2017 have shown that single cholinergic cells in the medial septum/ ventral diagonal band of Broca projected simultaneously to multiple brain areas, such as the mPFC and MeA. Collectively, our and other results indicate that cholinergic cells in the HDB and VP/SI predominantly send axons separately the BLA and mPFC, but a significant portion of ChAT+ neruons have dual projections towards both regions, an innervation strategy, which allows for simultaneous as well as independent control of operation in these cortical areas.

A portion of subcortical neuromodulatory afferents typically express more than one kind of neurotransmitter molecule. For instance, dopaminergic cells can release GABA or glutamate in addition to dopamine (Stuber et al., 2010; Tecuapetla et al., 2010; Tritsch et al., 2012, 2014). Similarly, BF cholinergic cells projecting to the neocortex and hippocampus typically release GABA as well (Saunders et al., 2015; Takács et al., 2018). Here, we showed that 70% of VP/SI cholinergic neurons co-express VGLUT3, while this ratio is ∼30% in the HDB. Moreover, VGLUT3-containing neurons and cholinergic cells in the VP/SI share similar projection patterns, innervating mostly to the mPFC and BA. We also found that 60% of cholinergic axon terminals in the BA contain VGLUT3, whereas this ratio is insignificant in other areas surrounding the amygdala. These results are in agreement with data obtained previously in rats (Poulin et al., 2006), showing that one of the two parallel BF projections to the amygdala, the VP/SI to BA pathway express significantly more VGLUT3, and therefore, may release both glutamate and acetylcholine.

In addition, our data suggest that cholinergic cells in the HDB and VP/SI transiently express VGAT during development. This conclusion is based on the fact that i) the vast majority of ChAT+ neurons in the HDB and VP/SI also expressed the reporter protein ZsGreen1 under the control of VGAT promoter in offspring of mice crossed by VGAT-Cre and Ai6 and ii) neither the labeling of VGAT+ cell bodies in the BF, nor the VGAT+ axon terminals in the BA showed immunoreactivity for ChAT in adult mice. These results can be explained if VGAT gene is transiently switches on during development, resulting in expression of ZsGreen1, the lifetime of which protein significantly outlasts the VGAT expression in cholinergic cells. Our findings, thus, show that GABAergic phenotype of ChAT+ neurons innervating the BLA is transient in agreement with recent results (Granger et al., 2023).

In summary, our results indicate that there are two parallel cholinergic pathways that are in the position to control simultaneously or independently the mPFC and BLA. This organizational principle of cholinergic projections can allow the maximalization of computational support for coordinated functioning of innervated brain regions. The uncovered structural arrangement sheds light on logic underlying cholinergic control of cortical function and has implications for understanding the neural basis of emotional regulation.

## Authors Contribution

Bence Barabas: performed research, design research, wrote the paper

Zsofia Reeb: performed research

Orsolya I Papp: design research, performed research

Norbert Hajos: design research, wrote the paper

## Nomenclature

Aca: anterior commissure, anterior part
ACo: anterior cortical amygdaloid nucleus
Acp: anterior commissure, posterior part
AHi: amygdalohippocampal area
AHiAL: amygdalohippocampal area, anterolateral part
AHiPM: amygdalohippocampal area, posteromedial part
AI: agranular insular cortex
APir: amygdalopiriform transition area
BA: basal amygdala
BAa: basal amygdala, anterior part
BAp: basal amygdala, posterior part
BFCNs: basal forebrain cholinergic neurons
BLA: basolateral amygdala
BMA: basomedial amygdala
CeL: central amygdala, lateral division
CeM: central amygdala, medial division
Cg1: cingulate cortex, area 1
Cg2: cingulate cortex, area 2
ChAT: choline acetyltransferase
DEn: dorsal endopiriform nucleus
DI: dysgranular insular cortex
DP: dorsal peduncular cortex
dPFC: dorsal prefrontal cortex
HDB: nucleus of the horizontal limb of the diagonal band
IL: infralimbic cortex
IPACL: interstitial nucleus of the posterior limb of the anterior commissure, lateral part
IPACM: interstitial nucleus of the posterior limb of the anterior commissure, medial part
LA: lateral amygdala
LEnt: lateral entorhinal cortex
LH: lateral hypothalamic area
LO: lateral orbital cortex
LPO: lateral preoptic area
M1: primary motor cortex
M2: secondary motor cortex
MCPO: magnocellular preoptic nucleus
MeA: medial amygdala, anterior part
Mep: medial amygdala, posterior part
MO: medial orbital cortex
mPFC: medial prefrontal cortex
MS: medial septal nucleus
Pir: piriform cortex
PL: prelimbic cortex
PMCo: posteromedial cortical amygdaloid nucleus
PS: parastrial nucleus
S1: primary somatosensory cortex
SI: substantia innominata
VAChT: vesicular acetylcholine transporter
VGLUT3: vesicular glutamate transporter 3
VO: ventral orbital cortex
VP: ventral pallidum
vPFC: ventral prefrontal cortex

## Acknowledgements

Acknowledgements: We acknowledge financial support from the HUN-REN Hungarian Research Network, Hungarian Brain Research Program (2017-1.2.1-NKP-2017-00002) and National Research, Development and Innovation Office (K131893 and RRF-2.3.1-21-2022-00004). The authors are grateful to Éva Krizsán and Erzsébet Gregori for their excellent technical assistance. We also thank László Barna, the Nikon Microscopy Center at the Institute of Experimental Medicine, Nikon Austria GmbH, and Auro-Science Consulting, Ltd., for kindly providing microscopy support.

## Figure Legends

**Suppl. Fig. 1.**
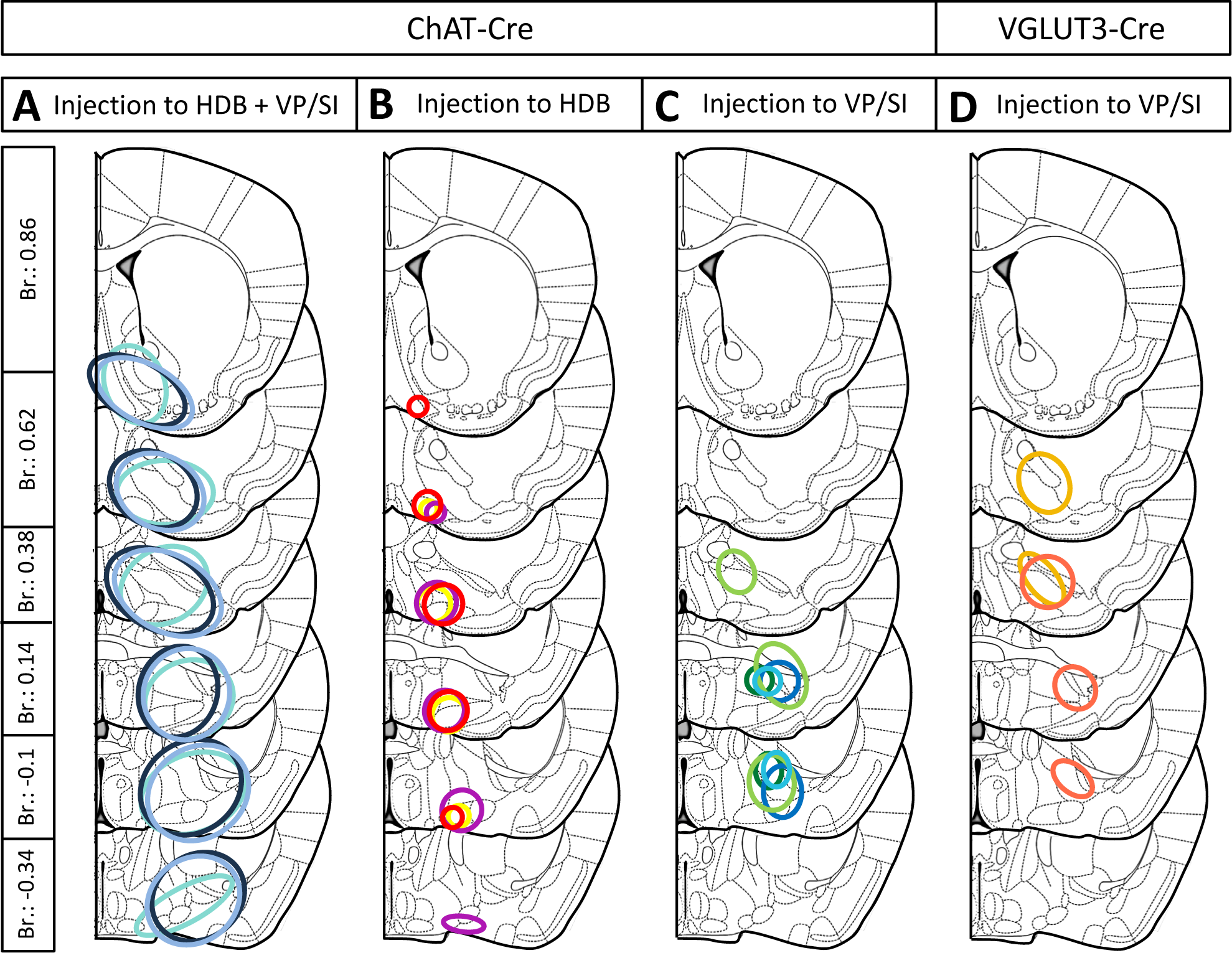
Reconstructed maps showing the localization and spread of virus injections used for BF anterograde labeling. (A) Localization and spread of 100+100nl AAV5.Ef1a.DIO.eYFP virus injection to the HDB and VP/SI in ChAT-Cre mice (n=3). Different colors represent different animals. These injections appertain to Figs 1 and 4. (B-C) Localization and spread of 30nl AAV8.CAG.flex.GFP virus injected to the HDB (B, n=3) or VP/SI (C, n=4) in ChAT-Cre mice. Different colors represent different animals. These injections appertain to Figs 3 and 5, and color codes are the same as in the corresponding figures. (D) Localization and spread of 30nl AAV8.CAG.flex.GFP virus in the VP/SI (n=2) in VGLUT3-Cre mice. Different colors represent different animals. These injections appertain to Fig 8.

**Suppl. Fig. 2.**
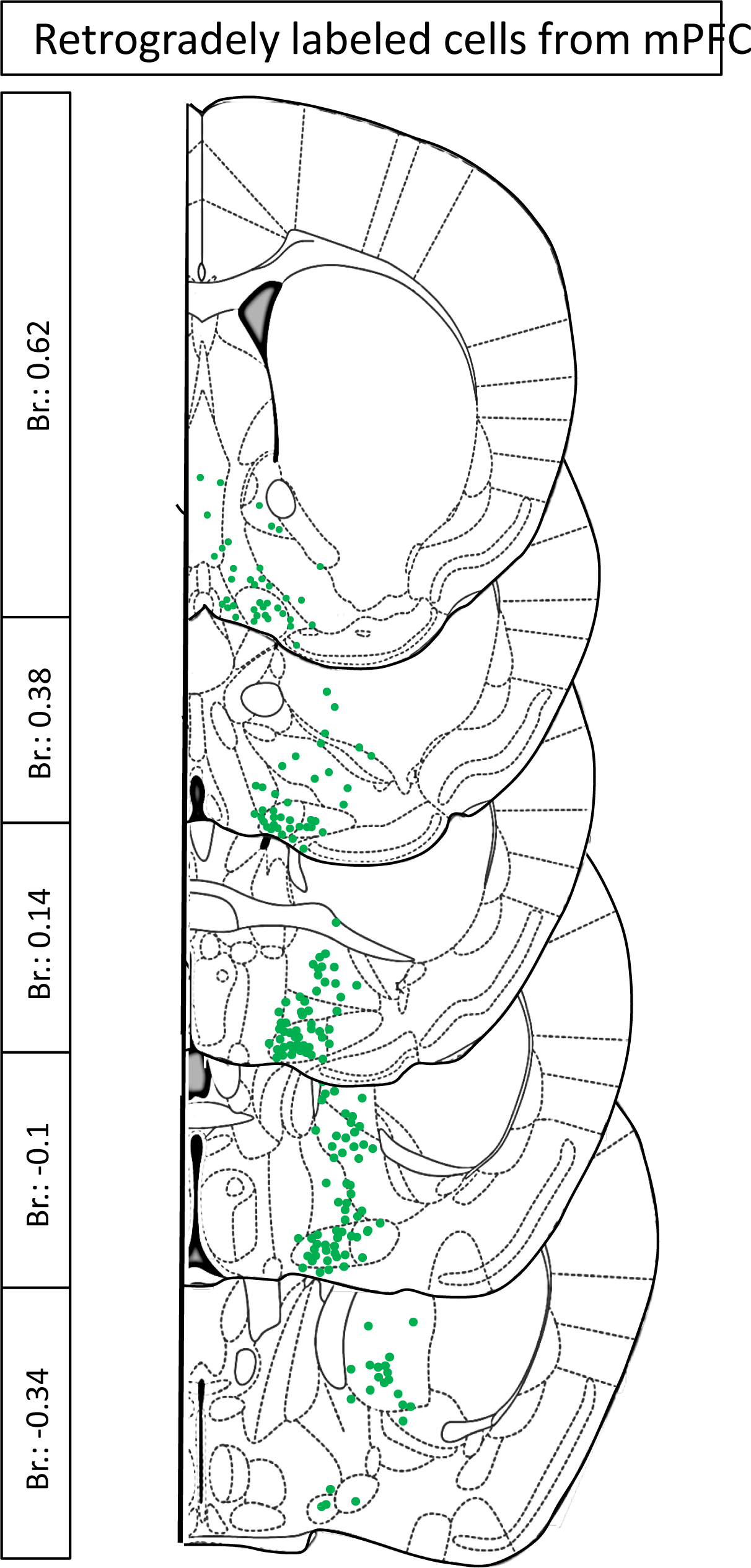
Localization of retrogradely labeled cholinergic BF neurons from retrograde-anterograde virus tracing. Green dots represent retrogradely labeled BF cholinergic neurons projecting to the mPFC. The area covered by greens dots defines the source of single and double-projecting cholinergic neurons in the BF.

**Suppl. Fig. 3.**
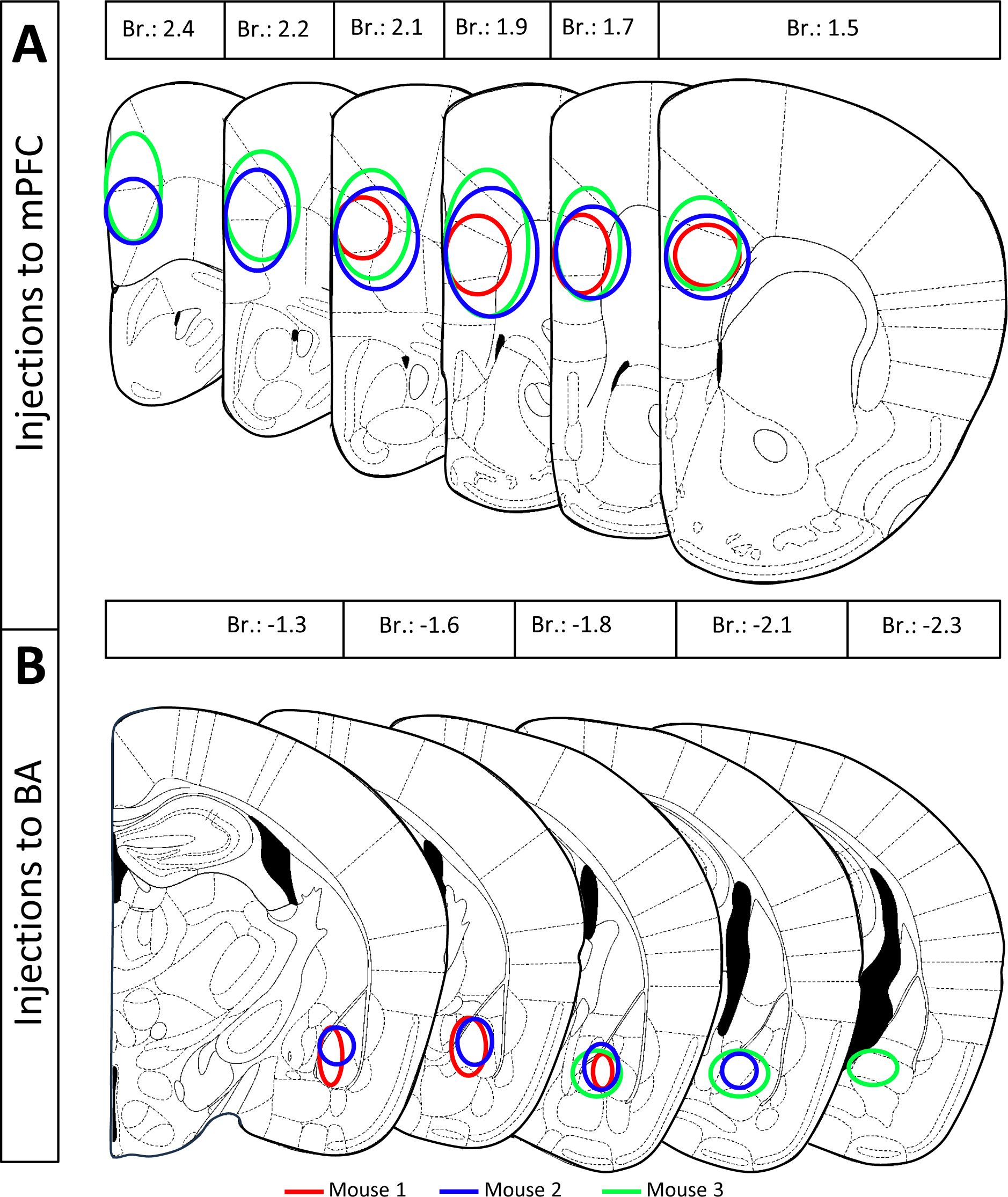
Reconstructed maps showing the localization and spread of retrograde tracers used for double-retrograde tracing. (A) Localization and spread of Fluorogold in the mPFC (n=3) (B) Localization and spread of Fast Blue in the BA (n=3). Different colors represent different animals.

**Suppl. Fig. 4.**
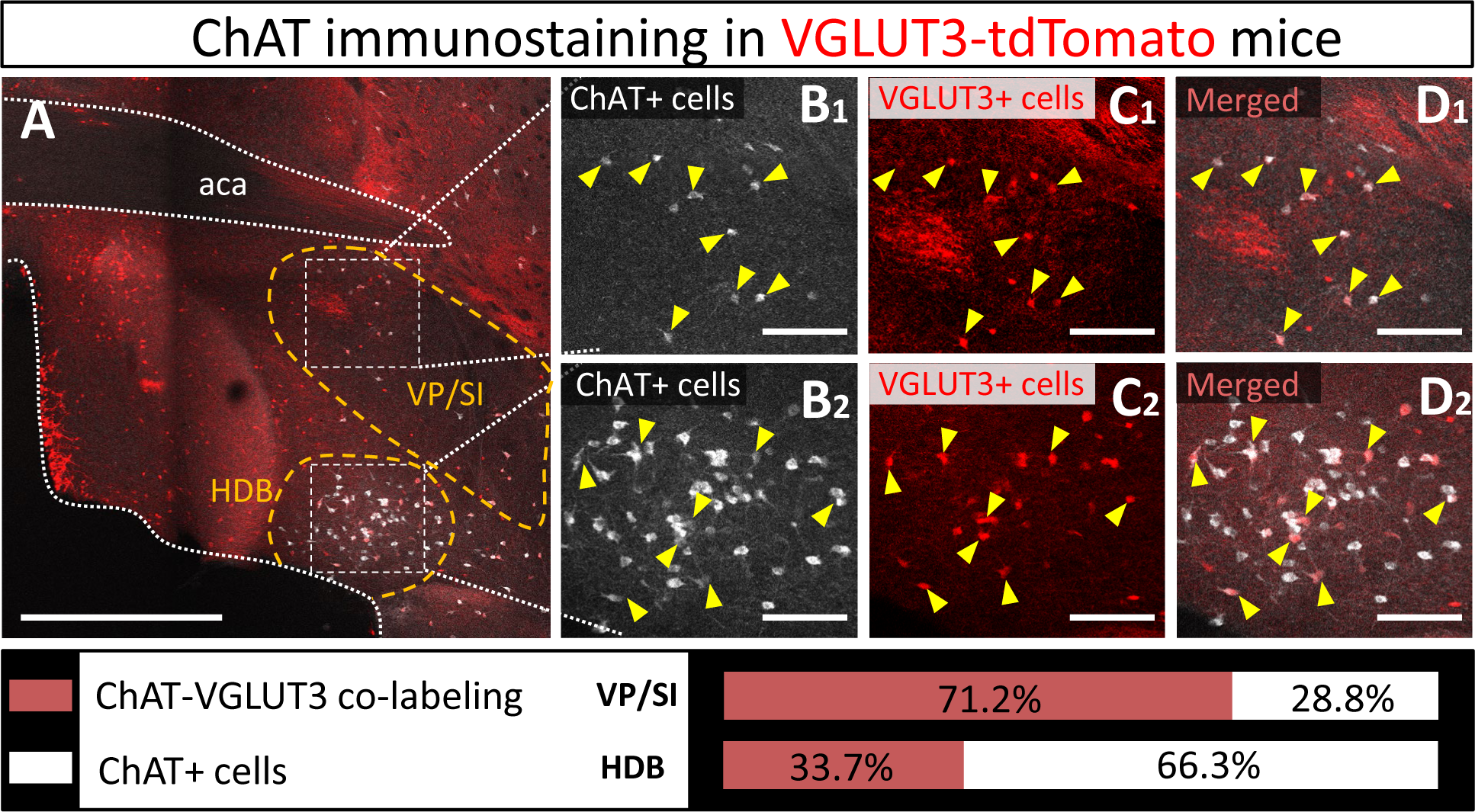
Ratio of ChAT and VGLUT3 co-labeling in the HDB and VP/SI. (A) Example coronal slice taken at the BF, showing ChAT content of VGLUT3-expressing neurons. The yellow dashed lines represent the borders of the HDB and VP/SI, where the co-labeling was counted. Scale bar: 500 µm. (B) ChAT+ cells in the VP/SI (B1) and HDB (B2). Scale bar: 100 µm (C) VGLUT3+ cells in the VP/SI (C_1_) and HDB (C_2_). Scale bar: 100 µm (D) Merged images taken at the VP/SI (D_1_) and HDB (D_2_). Scale bar: 100 µm. Yellow arrows indicate co-labeled neurons. Bottom graph showing the ratio of co-labeling in the VP/SI and HDB, respectively.

**Suppl. Fig 5.**
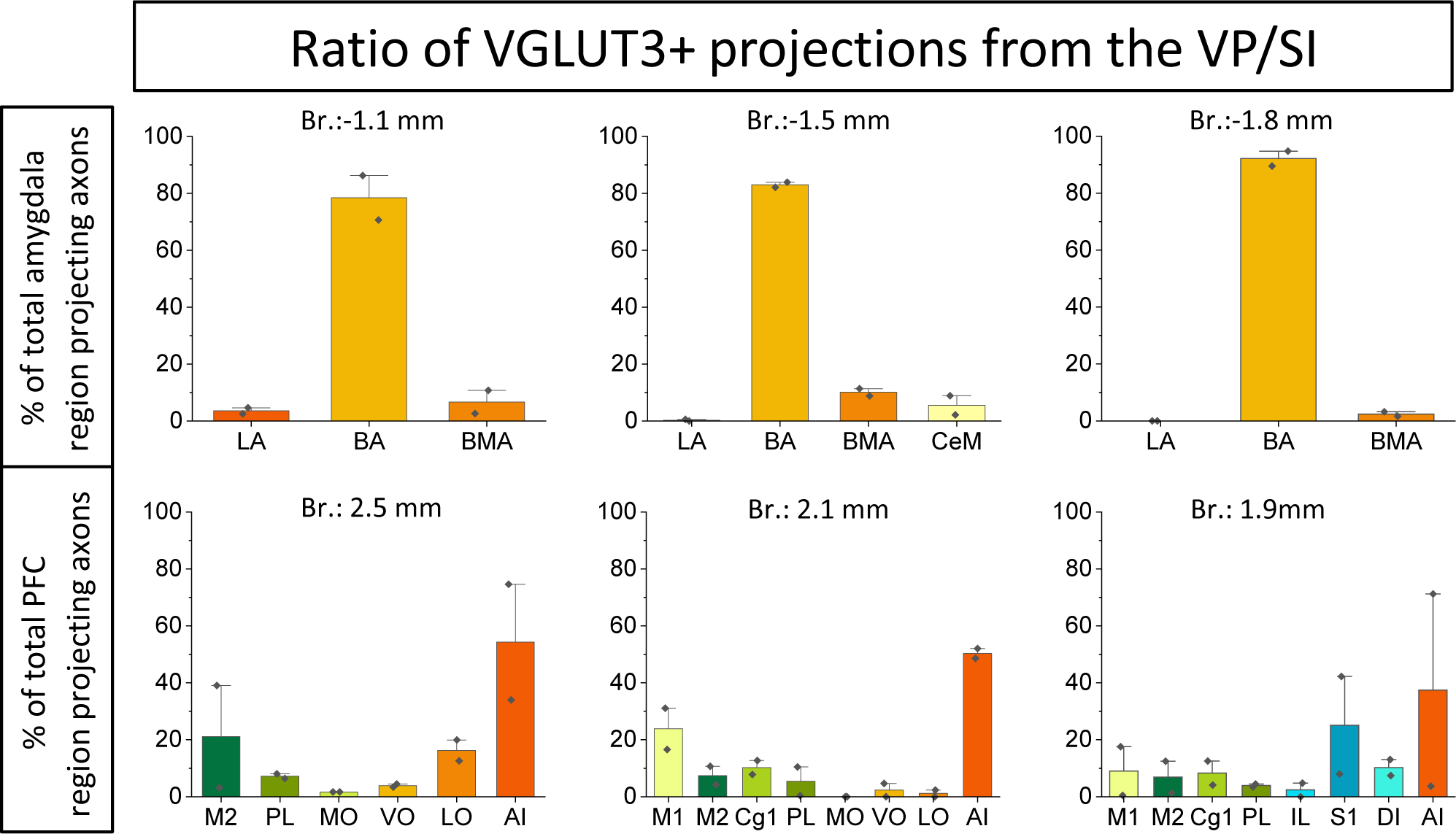
Ratio of glutamatergic (VGLUT3) projections in the amygdala and PFC subregions originating from the VP/SI. (A) Percentage of amygdala region-projecting VP/SI glutamatergic axons in each nucleus/region at different bregma levels. (B) Percentage of PFC region-projecting VP/SI glutamatergic axons in each nucleus/region at different bregma levels. These graphs appertain to Fig 8.

## References

1. Adhikari, A., Lerner, T. N., Finkelstein, J., Pak, S., Jennings, J. H., Davidson, T. J., Ferenczi, E., Gunaydin, L. A., Mirzabekov, J. J., Ye, L., Kim, S. Y., Lei, A., & Deisseroth, K. (2015). Basomedial amygdala mediates top-down control of anxiety and fear. Nature, 527(7577), 179–185. 10.1038/nature15698

2. Aitta-aho, T., Hay, Y. A., Phillips, B. U., Saksida, L. M., Bussey, T. J., Paulsen, O., & Apergis-Schoute, J. (2018). Basal forebrain and brainstem cholinergic neurons differentially impact amygdala circuits and learning-related behavior. Current Biology, 28(16), 2557–2569.e4. 10.1016/j.cub.2018.06.064

3. Bloem, B., Schoppink, L., Rotaru, D. C., Faiz, A., Hendriks, P., Mansvelder, H. D., van de Berg, W. D. J., & Wouterlood, F. G. (2014). Topographic mapping between basal forebrain cholinergic neurons and the medial prefrontal cortex in mice. Journal of Neuroscience, 34(49), 16234–16246. 10.1523/JNEUROSCI.3011-14.2014

4. Bloodgood, D. W., Sugam, J. A., Holmes, A., & Kash, T. L. (2018). Fear extinction requires infralimbic cortex projections to the basolateral amygdala. Translational Psychiatry, 8(1). 10.1038/s41398-018-0106-x

5. Burgos-Robles, A., Vidal-Gonzalez, I., & Quirk, G. J. (2009). Sustained conditioned responses in prelimbic prefrontal neurons are correlated with fear expression and extinction failure. Journal of Neuroscience, 29(26), 8474–8482. 10.1523/JNEUROSCI.0378-09.2009

6. Carlsen, J., Záborszky, L., & Heimer, L. (1985). Cholinergic projections from the basal forebrain to the basolateral amygdaloid complex: A combined retrograde fluorescent and immunohistochemical study. Journal of Comparative Neurology, 234(2), 155–167. 10.1002/cne.902340203

7. Cassell MD, Wright DJ. Topography of projections from the medial prefrontal cortex to the amygdala in the rat. Brain Res Bull. 1986 Sep;17(3):321–33. doi: 10.1016/0361-9230(86)90237-6. PMID: 2429740.

8. Crimmins, B. E., Lingawi, N. W., Chieng, B. C., Leung, B. K., Maren, S., & Laurent, V. (2023). Basal forebrain cholinergic signaling in the basolateral amygdala promotes strength and durability of fear memories. Neuropsychopharmacology, 48(4), 605–614. 10.1038/s41386-022-01427-w

9. Crouse, R. B., Kim, K., Batchelor, H. M., Girardi, E. M., Kamaletdinova, R., Chan, J., Rajebhosale, P., Pittenger, S. T., Role, L. W., Talmage, D. A., Jing, M., Li, Y., Gao, X. B., Mineur, Y. S., & Picciotto, M. R. (2020). Acetylcholine is released in the basolateral amygdala in response to predictors of reward and enhances the learning of cue-reward contingency. ELife, 9, 1–31. 10.7554/ELIFE.57335

10. Davis M. Neurobiology of fear responses: the role of the amygdala. J Neuropsychiatry Clin Neurosci. 1997 Summer; 9(3):382–402. doi: 10.1176/jnp.9.3.382. PMID: 9276841.

11. Farr SA, Uezu K, Flood JF, Morley JE. Septo-hippocampal drug interactions in post-trial memory processing. Brain Res. 1999 Nov 20;847(2):221–30. doi: 10.1016/s0006-8993(99)02049-1. PMID: 10575091.

12. Gabbott, P. L. A., Warner, T. A., Jays, P. R. L., Salway, P., & Busby, S. J. (2005). Prefrontal cortex in the rat: Projections to subcortical autonomic, motor, and limbic centers. Journal of Comparative Neurology, 492(2), 145–177. 10.1002/cne.20738

13. Goldbach, R., Allgaier, C., Heimrich, B., & Jackisch, R. (1998). Postnatal development ofmuscarinic autoreceptors modulating acetylcholine release in the septohippocampal cholinergic system I. Axon terminal region: hippocampus. In Developmental BrainResearch (Vol. 108). 10.1016/s0165-3806(98)00026-1.

14. Gombkoto, P., Gielow, M., Varsanyi, P., Chavez, C., & Zaborszky, L. (2021). Contribution of the basal forebrain to corticocortical network interactions. Brain Structure and Function, 226(6), 1803–1821. 10.1007/s00429-021-02290-z

15. Granger, A. J., Mao, K., Saulnier, J. L., Hines, M. E., & Sabatini, B. L. (2023). Developmental regulation of GABAergic gene expression in forebrain cholinergic neurons. Frontiers in Neural Circuits, 17. 10.3389/fncir.2023.1125071

16. Hasselmo, M. E., & Sarter, M. (2011). Modes and models of forebrain cholinergic neuromodulation of cognition. In Neuropsychopharmacology (Vol. 36, Issue 1, pp. 52–73). 10.1038/npp.2010.104

17. Herry, C., Ciocchi, S., Senn, V., Demmou, L., Müller, C., & Lüthi, A. (2008). Switching on and off fear by distinct neuronal circuits. Nature, 454(7204), 600–606. 10.1038/nature07166

18. Hurley, K. M., Herbert, H., Moga, M. M., & Saper, C. B. (1991). Efferent projections of the infralimbic cortex of the rat. Journal of Comparative Neurology, 308(2), 249–276. 10.1002/cne.903080210

19. Izquierdo, I., Furini, C. R. G., & Myskiw, J. C. (2016). Fear memory. Physiol Rev, 96, 695–750. 10.1152/physrev.00018.2015.-Fear

20. Jiang, L., Kundu, S., Lederman, J. D. D., López-Hernández, G. Y. Y., Ballinger, E. C. C., Wang, S., Talmage, D. A. A., & Role, L. W. W. (2016). Cholinergic signaling controls conditioned fear behaviors and enhances plasticity of cortical-amygdala circuits. Neuron, 90(5), 1057– 1070. 10.1016/j.neuron.2016.04.028

21. Jope RS. High affinity choline transport and acetylCoA production in brain and their roles in the regulation of acetylcholine synthesis. Brain Res. 1979 Dec;180(3):313–44. doi: 10.1016/0165-0173(79)90009-2. PMID: 394816.

22. Kim, J., Pignatelli, M., Xu, S., Itohara, S., & Tonegawa, S. (2016). Antagonistic negative and positive neurons of the basolateral amygdala. Nature Neuroscience, 19(12), 1636–1646. 10.1038/nn.4414

23. Kitt, C. A., Mitchell, S. J., Delong, M. R., Wainer, B. H., & Price, D. L. (1987). Fiber pathways of basal forebrain cholinergic neurons in monkeys. In Brain Research (Vol. 406, Issue 16). 10.1016/0006-8993(87)90783-9.

24. Knox, D. (2016). The role of basal forebrain cholinergic neurons in fear and extinction memory. In Neurobiology of Learning and Memory (Vol. 133, pp. 39–52). Academic Press Inc. 10.1016/j.nlm.2016.06.001

25. Lacroix, L., Spinelli, S., Heidbreder, C. A., & Feldon, J. (2000). Differential Role of the Medial and Lateral Prefrontal Cortices in Fear and Anxiety. Behavioral Neuroscience, 114(6), 1119–1130. 10.1037/0735-7044.114.6.1119

26. LeDoux JE. (2000). Emotion circuits in the brain. Annu Rev Neurosci. 2000; 23:155–84. 10.1146/annurev.neuro.23.1.155

27. Lee, S., & Kim, J. H. (2019). Basal forebrain cholinergic-induced activation of cholecystokinin inhibitory neurons in the basolateral amygdala. Experimental Neurobiology, 28(3), 320–328. 10.5607/en.2019.28.3.320

28. Li, X., Yu, B., Sun, Q., Zhang, Y., Ren, M., Zhang, X., Li, A., Yuan, J., Madisen, L., Luo, Q., Zeng, H., Gong, H., & Qiu, Z. (2017). Generation of a whole-brain atlas for the cholinergic system and mesoscopic projectome analysis of basal forebrain cholinergic neurons. Proceedings of the National Academy of Sciences of the United States of America, 115(2), 415–420. 10.1073/pnas.1703601115

29. Lingawi, N. W., Laurent, V., Westbrook, R. F., & Holmes, N. M. (2019). The role of the basolateral amygdala and infralimbic cortex in (re)learning extinction. In Psychopharmacology (Vol. 236, Issue 1, pp. 303–312). Springer Verlag. 10.1007/s00213-018-4957-x

30. Little, J. P., & Carter, A. G. (2013). Synaptic mechanisms underlying strong reciprocal connectivity between the medial prefrontal cortex and basolateral amygdala. Journal of Neuroscience, 33(39), 15333–15342. 10.1523/JNEUROSCI.2385-13.2013

31. Lozovaya, N., Eftekhari, S., Cloarec, R., Gouty-Colomer, L. A., Dufour, A., Riffault, B., Billon-Grand, M., Pons-Bennaceur, A., Oumar, N., Burnashev, N., Ben-Ari, Y., & Hammond, C. (2018). GABAergic inhibition in dual-transmission cholinergic and GABAergic striatal interneurons is abolished in Parkinson disease. Nature Communications, 9(1). 10.1038/s41467-018-03802-y

32. Mallet, P. E., Beninger, R. J., Flesher, S. N., Jhamandas, -khem, & Roland Boegman, –and J. (1995). Nucleus Basalis Lesions: Implication of Basoamygdaloid Cholinergic Pathways in Memory. In Brain Research Bulletin (Vol. 36, Issue I). 10.1016/0361-9230(94)00162-t.

33. Maren, S., Aharonov, G., Stote, D. L., & Fanselow, M. S. (1996). 7v-methyl-d-aspartate receptors in the basolateral amygdala are required for both acquisition and expression of conditional fear in rats. in behavioral neuroscience (Vol. 110, Issue 6). 10.1037//0735-7044.110.6.1365.

34. Mayo, W., Dubois, B., Ploska, A., Javoy-Agid, F., Agid, Y., Le Moal, M., & Simon, H. (1984). Cortical cholinergic projections from the basal forebrain of the rat, with special reference to the prefrontal cortex innervation. In Neuroscience Letters (Vol. 47). 10.1016/0304-3940(84)90421-x.

35. Mcdonald AJ, Mascagni F, Guo L. Projections of the medial and lateral prefrontal cortices to the amygdala: a Phaseolus vulgaris leucoagglutinin study in the rat. Neuroscience. 1996 Mar;71(1):55–75. doi: 10.1016/0306-4522(95)00417-3. PMID: 8834392.

36. Mesulam, M. –Marsel, Mufson, E. J., Levey, A. I., & Wainer, B. H. (1983). Cholinergic innervation of cortex by the basal forebrain: Cytochemistry and cortical connections of the septal area, diagonal band nuclei, nucleus basalis (Substantia innominata), and hypothalamus in the rhesus monkey. Journal of Comparative Neurology, 214(2), 170–197. 10.1002/cne.902140206

37. Mineur, Y. S., Mose, T. N., Maibom, K. L., Pittenger, S. T., Soares, A. R., Wu, H., Taylor, S. R., Huang, Y., & Picciotto, M. R. (2022). ACh signaling modulates activity of the GABAergic signaling network in the basolateral amygdala and behavior in stress-relevant paradigms. Molecular Psychiatry, 27(12), 4918–4927. 10.1038/s41380-022-01749-7

38. Nagai, T., Kimura, H., Maeda, T., Mcgeer, P. L., Peng, F., & Mcgeer2, E. G. (1982). Cholinergic projections from the basal forebrain of rat to the amygdala’. In Society for Neuroscience Printed in U.S.A (Vol. 2, Issue 4). 10.1523/JNEUROSCI.02-04-00513.1982

39. Okada, K., Nishizawa, K., Kobayashi, T., Sakata, S., Hashimoto, K., & Kobayashi, K. (2021). Different cholinergic cell groups in the basal forebrain regulate social interaction and social recognition memory. Scientific Reports, 11(1). 10.1038/s41598-021-93045-7

40. Pascual, M., Pérez-Sust, P., & Soriano, E. (2004). The GABAergic septohippocampal pathway in control and reeler mice: Target specificity and termination onto reelin-expressing interneurons. Molecular and Cellular Neuroscience, 25(4), 679–691. 10.1016/j.mcn.2003.12.009

41. Paxinos, G., Franklin, K. B. J., Paxinos, G and Franklin, K. B. J., Paxinos, G., & Franklin, K. B. J. (2004). Mouse Brain in Stereotaxic Coordinates. In Academic Press (Vol. 2nd). 10.1016/S0306-4530(03)00088-X

42. Petrovich, G. D., Risold, P. Y., & Swanson, L. W. (1996). Organization of projections from the basomedial nucleus of the amygdala: A PHAL study in the rat. Journal of Comparative Neurology, 374(3), 387–420. 10.1002/(SICI)1096-9861(19961021)374:3<387::AID-CNE6>3.0.CO;2-Y

43. Poulin, A. N., Guerci, A., Mestikawy, S. El, & Semba, K. (2006). Vesicular glutamate transporter 3 immunoreactivity is present in cholinergic basal forebrain neurons projecting to the basolateral amygdala in rat. The Journal Of Comparative Neurology, 498(5), 690– 711. 10.1002/cne.21081

44. Power, A. E., Vazdarjanova, A., & McGaugh, J. L. (2003). Muscarinic cholinergic influences in memory consolidation. Neurobiology of Learning and Memory, 80(3), 178–193. 10.1016/S1074-7427(03)00086-8

45. Robert, B., Kimchi, E. Y., Watanabe, Y., Chakoma, T., Jing, M., Li, Y., & Polley, D. B. (2021). A functional topography within the cholinergic basal forebrain for encoding sensory cues and behavioral reinforcement outcomes. Elife, 10:e69514. 10.7554/eLife

46. Santini, E., Quirk, G. J., & Porter, J. T. (2008). Fear conditioning and extinction differentially modify the intrinsic excitability of infralimbic neurons. Journal of Neuroscience, 28(15), 4028–4036. 10.1523/JNEUROSCI.2623-07.2008

47. Saunders, A., Granger, A. J., & Sabatini, B. L. (2015). Corelease of acetylcholine and GABA from cholinergic forebrain neurons. Elife, 4:e06412. 10.7554/eLife.06412.001

48. Senn, V., Wolff, S. B. E., Herry, C., Grenier, F., Ehrlich, I., Grundemann, J., Fadok, J. P., Muller, C., Letzkus, J. J., & Luthi, A. (2014). Long-range connectivity defines behavioral specificity of amygdala neurons. Neuron, 81(2), 428–437. 10.1016/j.neuron.2013.11.006

49. Sierra-Mercado, D., Padilla-Coreano, N., & Quirk, G. J. (2011). Dissociable roles of prelimbic and infralimbic cortices, ventral hippocampus, and basolateral amygdala in the expression and extinction of conditioned fear. Neuropsychopharmacology, 36(2), 529–538. 10.1038/npp.2010.184

50. Solari, N., & Hangya, B. (2018). Cholinergic modulation of spatial learning, memory and navigation. In European Journal of Neuroscience (Vol. 48, Issue 5, pp. 2199–2230). Blackwell Publishing Ltd. 10.1111/ejn.14089

51. Sotres-Bayon, F., Sierra-Mercado, D., Pardilla-Delgado, E., & Quirk, G. J. (2012). Gating of fear in prelimbic cortex by hippocampal and amygdala inputs. Neuron, 76(4), 804–812. 10.1016/j.neuron.2012.09.028

52. Stuber, G. D., Hnasko, T. S., Britt, J. P., Edwards, R. H., & Bonci, A. (2010). Dopaminergic terminals in the nucleus accumbens but not the dorsal striatum corelease glutamate. Journal of Neuroscience, 30(24), 8229–8233. 10.1523/JNEUROSCI.1754-10.2010

53. Takács, V. T., Cserép, C., Schlingloff, D., Pósfai, B., Szőnyi, A., Sos, K. E., Környei, Z., Dénes, Á., Gulyás, A. I., Freund, T. F., & Nyiri, G. (2018). Co-transmission of acetylcholine and GABA regulates hippocampal states. Nature Communications, 9(1). 10.1038/s41467-018-05136-1

54. Tecuapetla, F., Patel, J. C., Xenias, H., English, D., Tadros, I., Shah, F., Berlin, J., Deisseroth, K., Rice, M. E., Tepper, J. M., & Koos, T. (2010). Glutamatergic signaling by mesolimbic dopamine neurons in the nucleus accumbens. Journal of Neuroscience, 30(20), 7105–7110. 10.1523/JNEUROSCI.0265-10.2010

55. Tovote, P., Fadok, J. P., & Luthi, A. (2015). Neuronal circuits for fear and anxiety. Nat Rev Neurosci, 16(6), 317–331. 10.1038/nrn3945

56. Tritsch, N. X., Ding, J. B., & Sabatini, B. L. (2012). Dopaminergic neurons inhibit striatal output through non-canonical release of GABA. Nature, 490(7419), 262–266. 10.1038/nature11466

57. Tritsch, N. X., Oh, W. J., Gu, C., & Sabatini, B. L. (2014). Midbrain dopamine neurons sustain inhibitory transmission using plasma membrane uptake of GABA, not synthesis. ELife, 2014(3). 10.7554/eLife.01936

58. Tye, K. M., Prakash, R., Kim, S. Y., Fenno, L. E., Grosenick, L., Zarabi, H., Thompson, K. R., Gradinaru, V., Ramakrishnan, C., & Deisseroth, K. (2011). Amygdala circuitry mediating reversible and bidirectional control of anxiety. Nature, 471(7338), 358–362. 10.1038/nature09820

59. Vertes, R. P. (2004). Differential Projections of the Infralimbic and Prelimbic Cortex in the Rat. Synapse, 51(1), 32–58. 10.1002/syn.10279

60. Weiskrantz L. (1956). Behavioral changes associated with ablation of the amygdaloid complex in monkeys. J Comp Physiol Psychol. Aug;49(4):381–91. 10.1037/h0088009.

61. Zaborszky, L. (2002). The modular organization of brain systems. Basal forebrain: the last frontier. Progress in Brain Research, 136, 359–372. 10.1016/S0079-6123(02)36030-8

62. Zaborszky, L., Csordas, A., Mosca, K., Kim, J., Gielow, M. R., Vadasz, C., & Nadasdy, Z. (2015). Neurons in the basal forebrain project to the cortex in a complex topographic organization that reflects corticocortical connectivity patterns: An experimental study based on retrograde tracing and 3D reconstruction. Cerebral Cortex, 25(1), 118–137. 10.1093/cercor/bht210

63. Zaborszky L, Gaykema RP, Swanson DJ, Cullinan WE. Cortical input to the basal forebrain. Neuroscience. 1997 Aug;79(4):1051–78. doi: 10.1016/s0306-4522(97)00049-3. PMID: 9219967.

